# Vegetative stage and soil horizon determine direction and magnitude of rhizosphere priming effects in contrasting treeline soils

**DOI:** 10.1101/2024.03.03.582814

**Authors:** Jennifer Michel, Sébastien Fontaine, Sandrine Revaillot, Catherine Piccon-Cochard, Jeanette Whitaker

## Abstract

1. Treelines in high latitudes and high altitudes are considered sentinels of global change. This manifests in accelerated encroachment of trees and shrubs and enhanced plant productivity, with currently unknown implications for the carbon balance of these biomes. Given the large soil organic carbon stocks in many treeline soils, we here wondered whether introducing highly productive plants would accelerate carbon cycling through rhizosphere priming effects and if certain soils would be more vulnerable to carbon loss from positive priming than others.
2. To test this, organic and mineral soils were sampled above and below treelines in the Swedish sub-arctic and the Peruvian Andes. A greenhouse experiment was then performed to quantify plant-induced changes in soil mineralisation rates (rhizosphere priming effect) and new C formation using natural abundance labelling and the C4-species Cynodon dactylon. Several environmental, plant, soil and microbial parameter were monitored during the experiment to complement the observations on soil C cycling.
3. Priming was predominantly positive at the beginning of the experiment, then systematically decreased in all soils during the plant growth season to be mostly negative at the end of the experiment at plant senescence. Independent of direction of priming, the magnitude of priming was always greater in organic than in corresponding mineral soils, which was best explained by the higher C contents of these soils. Integrated over the entire study period, the overall impact of priming (positive and negative) on the soil C balance was mostly negligible. Though, net soil C loss was observed in organic soils from the sub-arctic tundra in Sweden.
4. Most notably, positive and negative priming effects were not mutually exclusive, rather omnipresent across ecosystems, depending on sampling time. The direction of priming seems to be fluctuating with plant productivity, rhizosphere carbon inputs and nutrient uptake. This highlights the need for integrative long-term studies if we aim to understand priming effects at ecosystem scale and greenhouse and laboratory studies must be validated in situ to enable reliable ecological upscaling.

## 1 Introduction

Ecosystems in high altitudes and high latitudes share two features, which are of particular importance in the context of global change: their soils contain large carbon (C) stocks (Zimmermann et al. 2010; Saatchi et al. 2011; Rolando et al. 2017; Yang et al. 2018) and these biomes are predicted to experience greater than average warming (Wookey et al. 2009; Cox et al. 2013; Classen et al. 2015; IPCC 2021). Several studies show that current and future warming directly impacts plant-soil interactions in these regions (KeuperWild et al. 2020; Nottingham et al. 2020, Garcia-Palacios 2021). Plants directly transfer photosynthetically fixed C to soil microbes via exudation, and plant tissue turnover provides further organic inputs. Soil microbes feed on these substances and lead the processing of labile C inputs, which can be transferred into biomass or invested into nutrient acquisition from soil organic matter (SOM) (Kuzyakov & Domanski 2000; Kuzyakov & Cheng 2001; Jones et al. 2009; Kaiser et al. 2010, 2011; Guyonnet, et al. 2018; Canarini et al. 2019). Organic inputs to soils increase with warming-induced increased plant productivity, but increased organic inputs are not directly proportional to soil C storage, because organic inputs can enhance soil organic matter (SOM) mineralization by microbes and thus increase the amount of CO_2_ released to the atmosphere. This phenomenon of altered mineralisation of SOM through modified microbial activity is known as a priming effect (“PE”, Löhnis 1926; Bingemann et al. 1953 but see Kuzyakov et al. 2000). Priming effects can either increase soil C mineralisation (positive priming effect) or reduce rates of SOM-mineralisation (negative priming effect).

Priming effects describe the change in soil mineralisation rates caused by labile organic inputs, while rhizosphere priming effects (RPE) specifically describe changes in SOM mineralisation rates induced by plant root activities such as rhizodeposition (Kuzyakov, 2002). Often, studies simulate rhizodeposition in controlled laboratory experiments adding isotopically labelled substrates to soils. This approach provides mechanistic insights about priming effects driven by microbes. However, soil incubations neglect factors such as diurnal and seasonal fluctuations in plant metabolic activities, including plant nutrient uptake and root elongation and exudation, which also influence soil carbon cycling. Soil incubations, therefore, have limited ecological relevance and can even be misleading as they merely capture a snapshot of continuous plant-soil-microbe interactions. The lack of comprehensive RPE studies is mainly caused by the practical challenges and often high costs of studying RPEs *in situ* and *in vivo* (Kuzyakov, 2002; Cros et al. 2019). For a full picture, continuous studies are however urgently needed, because C inputs by plants are continuously changing in quantity and quality, which is also the case for plant nutrient uptake, and consequently RPE cannot be understood nor upscaled from single-time measurements and reductionist laboratory experiments without plants. One approach to capture rhizosphere processes at the mesocosm level is to take advantage of the natural difference of fractionation between heavy and light carbon isotopes (12-C to 13-C) in C_4_-plants, introducing them to C_3_ soils (Balesdent et al., 1987; Martin et al. 1990). This natural abundance labelling approach provides continuous and realistic carbon inputs with a traceable 13-C signature, which allows to study C cycling between plants, soils and microbes over relevant time scales, without modifying nutrient cycles. Eventually, the isotopic ratio of plant C inputs can be used to partition old (C_3_-soil) and new (C_4_-plant) C in soil respiration and microbial biomass C. In this study, natural abundance labelling was used to test in how far the carbon stocks of high altitudinal and high latitudinal treeline soils would be mobilized when highly productive plants are introduced. The specific focus was on differences in RPE between soil origin (boreal subarctic, Andean tropics), land cover types (boreal or tropical forest, tundra heath, Puna grassland) and soil horizons (organic top soils, mineral sub-soils), to answer the question which of these regions would be most vulnerable to soil C losses from priming.

## 2 Material and methods

### 2.1 Soil sampling

Soils were collected in 2016 in the high altitudes of the Peruvian Andes in Manú National Park in the department of Cusco at an average elevation of 3300 m (13°07’S 71°36’W, sampling May - June) and in the high latitudes of the boreal subarctic near the Abisko Scientific Research Station, 250 km north of the Arctic Circle in Northern Sweden at an average elevation of 650 m (68°21’N 18°49’E, August-September). The study area in the Peruvian Andes is situated at the high end of the Kosñipata transect on the Eastern side of the Andes, on the Western-facing hill side of the Paucartambo river valley. The study area comprises a montane tropical forest with a short transition zone leading into Puna grassland. The forest is a high Andean tropical mountain forest dominated by *Weinmannia microphylla* (Kunth), *Polylepis pauta* (Hieron.) and Gynoxys induta (Cuatrec.). The adjacent Puna grasslands are mainly composed of the genera *Festuca, Hypericum* and *Carex*. The climate is characterised by a rainy season from October to April, but in the forest and at the treeline cloud cover can be dense and humidity high throughout the year. The mean temperature is around 13 °C at the treeline, but can reach up to 25 °C in October and cool down to 3–6 °C in the Puna (UNEP World Conservation Monitoring Centre 2017). The soils referred to as “Andean soils” in this study are derived from volcanic material with mostly low base status. Because of diverse topography, slope and exposure, and due to varying history of erosion and landslides, these soils represent a variety of soil types. The forest soils are mostly Cambisols with a large fraction of organic matter in the upper soil horizon. The Puna grassland soils are shallower and mostly Andosols, where the sub soil contains notable quantities of amorphous clay (FAO Soil map of the world 1971; Wilcox et al. 1988; FAO World reference base for soil resources 2015). The study region in the Swedish subarctic is located near Abisko, south of the lake Torneträsk. The treeline transition along the elevational gradient has a Northeast—Southwest orientation. The studied treeline forms the upper end of a fragmented birch forest, which fades into alpine tundra. The dominant canopy-forming species of the studied birch forests is *Betula pubescens* (Ehrh.), while at some sites *Betula nana* (L.), *Salix glauca* (L.) and *Juniper* sp. are also present. The forest understorey is mostly composed of ericaceous plants such as *Empetrum nigrum* (L.), and several species of *Vaccinium*. The plant species composition of the upland heath lands is similar to the forest understorey, mainly composed of dwarf shrubs and cryptogams. In contrast to the Andean uplands, true grasses are widely absent, but species of the genera *Lycopodium* and *Equisetum* are commonly present at low abundance. There is regularly snow on the ground until late May and while average temperatures may be a little over 10 °C in July, by mid-August the average temperature is already declining rapidly with frosts likely by early September. In winter, temperatures can drop down to—34 °C. Precipitation averages 15 mm per month during the year, with July and August being wetter (60 mm/month; Abisko Scientific Research Station). Bedrock is formed by salic igneous rocks and quartic and phyllitic hard schists (Sundqvist et al. 2011). The subarctic soils referred to as “Boreal soils” in this study are permafrost-free and mostly Podsols and Cambisols with thin organic rich topsoils and sandy mineral soils from the B-horizon (FAO World reference base for soil resources 2015).

Eight soil types were classified representing the treeline ecotone based on the soil origin in terms of geographic region (Andean, Boreal), native current land cover (tropical mountain Forest or boreal Forest, Puna grassland, Tundra heath) and soil horizon (Organic, Mineral). We follow the same labelling throughout the manuscript, defining the soils by these three characteristics as: Andean Forest Organic (AFO), Andean Forest Mineral (AFM), Andean Puna organic (APO), Andean Puna Mineral (APM), Boreal Forest Organic (BFO), Boreal Forest Mineral (BFM), Boreal Tundra Organic (BTO) and Boreal Tundra Mineral (BTM). From both regions, fresh undisturbed samples were kept in sealed plastic bags in cool boxes at 4°C until the beginning of the experiment in early 2018. Before the experimental phase in the greenhouse, individual soil samples were carefully homogenized by hand, remaining large roots and rocks were removed and a sub-sample of each soil type was analysed for micronutrient contents using calcium-acetate-lactate (CAL) extraction (phosphorous, potassium), calcium chloride (magnesium) or calcium chloride/DTPA (CAT) extraction (sodium, sulphate, copper, zinc, boron, manganese) followed by spectrometric detection (Agilent 5110 ICP-OES) and for soil texture (sand, silt, clay) via wet sieving (Table 1, supplementary Table S1).

**Table 1:**
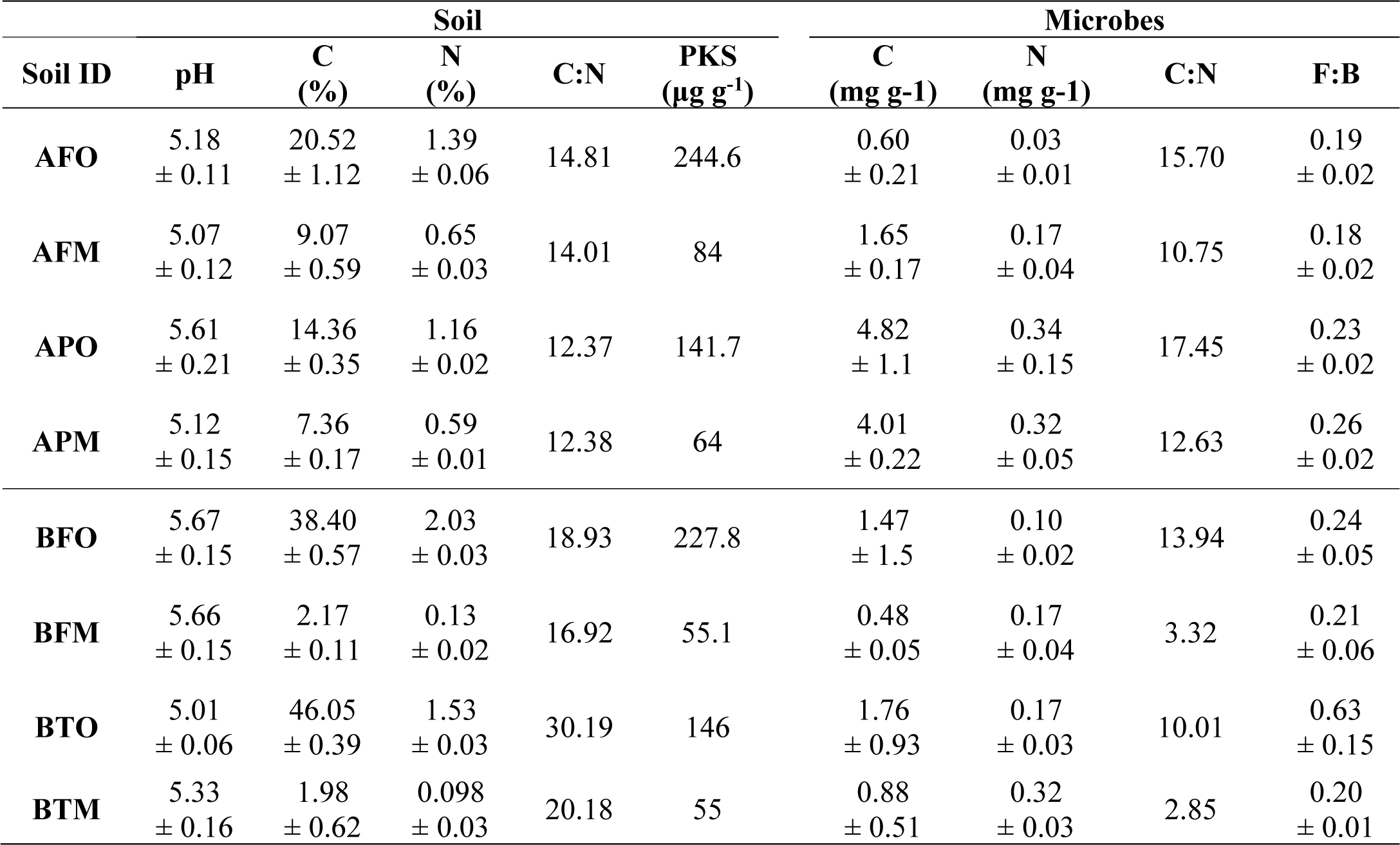
Key soil and microbial characteristics of the eight studied soil types. Abbreviations: C: carbon, N: nitrogen, PKS: phosphorous (P), potassium (K) and sulphur (S), F:B: fungal to bacteria ratio

### 2.2 Setting up the potting experiment in the greenhouse

For each soil type, eight pots (d = 5.5 cm, V = 1.02 l) were filled with equal amounts of soil per pot and a density between 0.4 g cm^-3^ and 0.15 g cm^-3^ depending on soil type (soil weight per pot at initial field moisture content: Peru mineral: 200 g, Peru organic: 150 g, Sweden mineral: 400 g, Sweden organic: 150 g). At the beginning of August 2018, four pots were randomly selected from each soil type and planted with the C_4_-species *Cynodon dactylon* (L.) Pers. at a density of 15 seeds per pot. *C. dactylon* is a fast-growing C_4_-grass, suitable for continuous cultivation in small mesocosms. Four pots of each soil type were maintained as unplanted controls (n=64 pots in total). For each soil type, maximum water holding capacity and permanent wilting point were determined via centrifugation (Revaillot et al. 2021). Six of the eight pots of each soil type (three planted and three unplanted) were equipped with soil moisture sensors (EC-5, Meter Group, Inc., Washington, USA), connected to a data logger (Campbell Scientific, Inc., Utah, USA). A commercial irrigation system (Rain Bird Corporation, California, USA) was modified to tune the water flow to a rate suitable to pot size (maximum of 10 ml per rain event) and adjusted to permanently maintain soil water content of all soil types (planted and unplanted) between the two levels of maximum water holding capacity and permanent wilting point, meaning around 30% volumetric soil moisture content (supplementary material S2). Pots were arranged in a segregated block design to match the tubing and valve system of the irrigation unit. The experiment was then run for three months from the sowing date (2^nd^ August 2018). The average greenhouse temperature during this period was 24±4°C (supplementary material S2), corresponding to 3-5°C warming of present-day summer temperatures for both Manú NP in Peru and Abisko in Sweden, which is in line with the RCP 8.5 scenario for the year 2100 (IPCC 2013; Abisko Scientific Research Station 2016; UNEP World Conservation Monitoring Centre 2017).

### 2.3 Soil respiration measurements

Over the course of three months of plant development, three respiration measurements were conducted to estimate RPE *in vivo*: RPE 1 was measured 35 days after sowing when plants were fully emerged, RPE 2 was measured 80 days after sowing when all plants had at least four leaves fully developed and RPE 3 was measured 95 days after sowing when the plants showed signs of senescence. For each measurement, the pots were removed from the greenhouse and each pot was placed in an individual dark respiration chamber (6.53l) under controlled temperature conditions (21.05 ± 1.15 °C). Each respiration chamber with a pot inside was flushed with CO_2_-free air for 45 seconds immediately before they were closed. Planted pots were then incubated for 24 hours whereas unplanted control pots containing only soil without plant were incubated for 48 hours to get sufficient concentrations of headspace CO_2_ for analyses. At the end of respective incubation period, from each chamber a 120 ml gas sample was taken in a glass flask sealed with air-lock rubber stoppers for subsequent analysis of CO_2_-concentration and isotopic composition. Samples were analysed on a gas chromatograph (Clarus 480, Perkin Elmer, Waltham, Massachusetts, United States) and a cavity ring down spectrometer (Picarro G2101i, Santa Clara, California, United States). All values were normalised for the time of respective incubation period. CO_2_-concentrations (ppm) were transformed to quantities of CO_2_-C (μg) according to

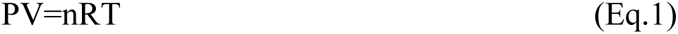

where P, V and T are the pressure (Pa), the volume (m3) and the temperature of the respiration chambers at the time of the gas sampling, respectively. The term n is the number of moles of C released by soil respiration and and R a constant (8.31 Jmol^-1^K^-1^).

### 2.4 Partitioning of carbon sources and quantification of RPE and new SOC formation

The difference of ^13^C isotopic composition between the C_3_ soils and the C_4_ plant (*Cynodon dactylon*) allowed us to partition total CO_2_-respiration into its soil and plant sources using following equations:

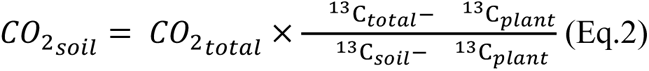

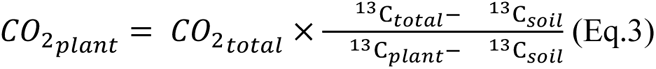

where CO_2_-_total_ and ^13^C_total_ are respectively the total CO_2_ flux and its ^13^C from plant-soil respiration; CO_2_-_soil_ and ^13^C_soil_ are respectively the CO_2_ flux and its ^13^C from microbial respiration of native (pre-existing) SOC and root litter, which have a C_3_-plant isotopic signature; and CO_2_-_plant_ and ^13^C_plant_ are respectively the CO_2_ flux and its ^13^C from respiration of the carbon fixed by the C_4_ species *Cynodon dactylon*. CO_2_-_plant_ includes *Cynodon dactylon* respiration as well as microbial respiration of C compounds deposited by this plant (litter, rhizodeposits).

At each measurement time, we used the average ^13^C from control soil respiration from all control pots of each soil type (n=4 per soil type) as value for the ^13^C of soil (^13^C_Soil_). To obtain accurate values for the ^13^C of plant inputs (^13^C_Plant_), we collected three leaves from each planted pot 24 hours after the pot incubations. The three leaves from each pot were placed into 120 ml glass flasks that were flushed with CO_2_-free air. After flushing, the flasks were sealed with rubber stoppers and incubated for 30 min at room temperature and in dark. The ^13^C of leaf respiration was measured using the same isotope laser analyser as for analysing the CO_2_ emitted from plant-soil systems (CRDS Analyser, Picarro, Santa Clara, CA, USA). At the end of the experiment, we repeated this analysis with root samples as well and compared the ^13^C of respiration of tissues of both leaves and roots to inform an uncertainty analysis (supplementary materials S3, S4).

### 2.5 Primed CO_2_ and SOC formation

Following the partitioning in section 1.4, the quantity of primed CO_2_-C can be estimated by comparing the soil-derived CO_2_-C in the planted pots and the CO_2_-C released in the unplanted control pots:

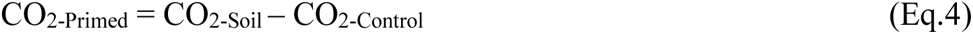

where CO_2-Soil_ refers to the CO_2_ from soil in the planted pots (Eq.2).

This provides a proxy of the rhizosphere priming effect, defined as the difference of CO_2_ released from SOM in presence of a plant compared to an unplanted control soil where all CO_2_ originates from SOM (Subke et al. 2006; Kuzyakov 2011). The amount of CO_2_-C primed was then expressed relative to the dry weight of soil and for a normalised incubation time of 24 hours for all samples (µg CO_2_-C g^-1^ soil day^-1^). At each time point, we used the average ^13^C from control soil respiration from all control pots of each soil type (n=4 per soil type) as value for the ^13^C of soil (^13^C_Soil_). To obtain accurate values for the ^13^C of plant inputs (^13^C_Plant_), we collected three leaves from each planted pot 24 hours after the pot incubations, placed them into 120 ml glass flasks, flushed the flasks with CO_2_-free air, sealed the flasks with rubber stoppers, incubated them for 30 min at room temperature and then measured the ^13^C of leaf respiration by directly connecting the flasks to the Picarro for 30 minutes. At the end of the experiment, we repeated this analysis with root samples as well and compared the ^13^C of respiration of tissues of both leaves and roots to inform an uncertainty analysis (supplementary Table S3, Fig. S4). An estimate for the total amount of primed C during the full course of the experiment (All prime) was generated by fitting 2^nd^ order polynomial regression lines through the three temporal measurements of each RPE for each sample and integrating the area under the graph between the first measurement of RPE (RPE1, 35 days after sowing) and the final RPE measurement (RPE3, 95 days after sowing):

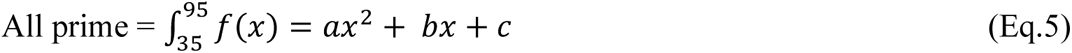

where x is the boundary for which we integrate (x=35 to x=95) to obtain the total amount of primed C during the 60 days corresponding to the measurement period. The y-axis in the integration has the unit (µg CO_2_-C g^-1^ soil), but we express the total amount of primed C for each soil type as (mg CO_2_-C g^-1^ soil) for improved comparability with values of new SOC formation and improved readability of the axis label.

New soil organic matter formation was calculated at the end of the experiment using the isotopic composition of bulk soil C in planted and unplanted soils and the ^13^C of root material:

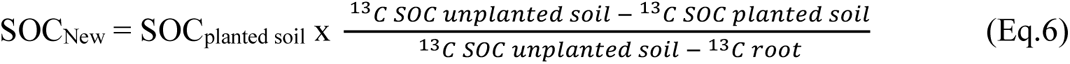

The SOC balance for this experiment was calculated as the difference between newly formed SOC (SOC_New,_ mg C g^-1^ soil) and the integral of all priming effects (All prime, mg CO_2_-C g^-1^ soil).

### 2.6 Plant parameters

Accompanying the first and second incubations in September and October (RPE1 and RPE2), several plant parameters were measured: Photosynthesis (PS) was measured at leaf level with three replicates per plant (n=12 per soil type) using a portable infrared gas analyser connected to a ¼ L chamber (LI-6200, Licor, Lincoln, Nebraska, United States). Corresponding leaf area was measured using a scanner (LI-3100C, Licor, Lincoln, Nebraska, United States) and photosynthetic active radiation (PAR) and leaf area index (LAI) of whole pots were measured using a ceptometer (Accupar LP-80, Meter Group, Pullman, Washington, United States), with one segment adapted to fit the size of the pots. Given the short time between the second and third RPE, these measurements were not repeated for the final RPE in November (RPE3).

### 2.7 Post-harvest biochemical analysis

At the end of the experiment, plant and soil material from each replicate pot (four planted and four unplanted controls per soil type) were separated and several parameters were measured. These comprised above and belowground biomass, ^12/13^C and N of plant tissues and bulk soil, soil mineral N, pH and moisture. For ^12/13^C and N analysis, plant and soil material was oven-dried (60 °C and 105 °C respectively), weighed, ground and analysed on an elemental analyser coupled to an isotope-ratio mass spectrometer (Vario isotope cube, Elementar, Langenselbold, Germany). Mineral N was extracted with 20 ml 1M KCl from subsamples of 5 g soil for each replicate. Extracts and blanks were analysed using a TOC/TN analyser (Shimadzu TNM-L, Shimadzu, Kyoto, Kyoto, Japan). Soil pH was measured on 5 g subsamples of soil in 12.5 ml deionised water (Schott instruments Lab870, Bath, Somerset, United Kingdom). Soil water content was determined by weight loss of 10 g fresh soil oven-dried at 105 °C for 48 hours.

Microbial biomass was extracted from 5 g soil using 30 mM K_2_SO_4_ and the vapour-phase chloroform fumigation technique modified after Vance et al. (1987). Extracts were lyophilised and elemental C and N contents and isotopic composition of the carbon compound were analysed on an elemental analyser coupled to an isotope-ratio mass spectrometer (Vario isotope cube, Elementar, Langenselbold, Germany). Microbial C and N were calculated by subtracting the respective amounts of non-fumigated (NF) extracts from the fumigated (F) ones. Given the heterogeneity of the soil types, no single correction factor was applied, and values are presented as they were measured following the recommendation in Halbritter et al. (2020), protocol 2.2.1 (Schmidt, I.K., Reinsch, S., Christiansen, C.T.).

Microbial biomass (MB) ^13^C was calculated as:

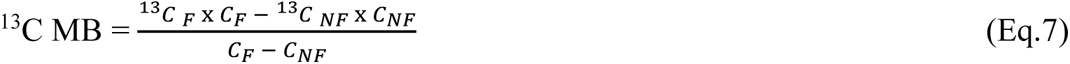

where ^13^C is the isotopic abundance and C is the amount of microbial biomass carbon per gram of dry weight soil (mg C g ^-1^ soil) as determined after salt-extraction in fumigated (F) and non-fumigated (NF) samples respectively.

The proportion of microbial biomass carbon which was soil-derived (MB_S_) was calculated as

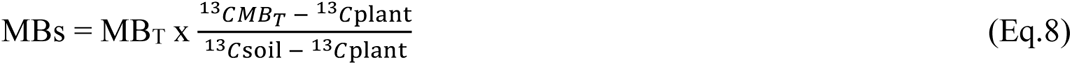

where MB_S_ is the soil-derived microbial biomass carbon (mg C g ^-1^ soil) and MB_T_ is the total microbial biomass carbon (mg C g ^-1^ soil), and ^13^C is the isotopic abundance of total microbial biomass C (MB_T_), plant C and soil C respectively.

The proportion of microbial biomass carbon which was plant-derived (MB_P_) was calculated by subtracting the soil-derived microbial biomass carbon from the total microbial biomass carbon (MB_T_):

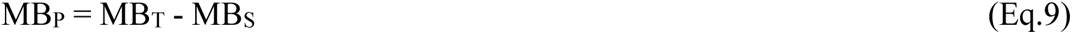

where MB_S_ is the soil-derived microbial biomass carbon (mg C g ^-1^ soil) and MB_T_ is the total microbial biomass carbon (mg C g ^-1^ soil) and MB_P_ is the plant-derived microbial biomass carbon (mg C g ^-1^ soil).

Microbial biomass and PLFAs have been characterized in control and planted pots both at the end of the experiment, to compare the two treatments planted and unplanted. Similar to the calculation of rhizosphere priming effects, which uses the unplanted control soils as reference, the microbial biomass of unplanted control soils was used as reference to describe plant-induced changes and is referred to as ‘initial’ microbial biomass, assuming only minor changes occur in microbial communities in unplanted control soils (Blagodatskaya et al. 2011; Lerch et al., 2011).

### 2.8 Phospholipid-derived fatty acids (PLFAs) of soil microorganisms

Microbial community composition of fungi, actinomycetes and gram-positive and gram-negative bacteria were determined by analysing phospholipid fatty acids (PLFAs) from both unplanted and planted soils at the end of the experiment. Subsamples of 10 g soil were frozen at -80°C and then freeze-dried and ground. Extraction was performed following a modified Bligh-Dyer protocol (White et al. 1979) and fatty acid methyl esters (FAMEs) were analysed on an Agilent 6890 gas chromatograph coupled to a mass spectrometer detector (Agilent 5973, Santa Clara, California, United States). Concentrations were blank corrected, normalised against an internal standard (methyl nonadecanoate C13:0 and C21:0, Sigma Aldrich, Merck Life Science UK Limited, Dorset, UK), classified by standard nomenclature (Frostegard et al. 1991, 1993) and reported on a soil mass basis as PLFA (µg g^-1^ dry weight soil). Biomarkers were assigned after Rinnan & Baath (2009), adding 18:1(n-9) for fungi (against Zelles 1997, but see Treonis et al. 2004; DeDeyn et al. 2011) and with the methyl-branched saturated fatty acids 10Me-16:0, 10Me-17:0 and 10Me-18:0 for actinomycetes after Zelles (1997, 1999). The two fungal biomarkers were tested for correlation (R^2^ between 0.88 and 0.96 in all batches) to avoid mis-ID-ing gram-bacteria through the 18:1(n-9) biomarker (Zelles, 1997). Final assignments were: saprophytic fungi (18:2, 18:1(n-9)), actinomycetes (10Me-16:0, 10Me-17:0, 10Me-18:0), other gram positive bacteria (15:0i, 15:0a, 16:0i, 17:0i, 17:0a), gram negative bacteria (16:1(n-7), cy17:0, 18:1(n-7), cy-19:0) and remaining unspecified PLFAs (14:0i, 14:0, 15:1i, 15:1a, 16:1(n-9), 16:1(n-5), 16:0, 17:1i(n-8), 17:0, 18:0i, 18:1(n-5), 18:0, 19:1). In this conservative assignment, the group of unspecified PLFAs also comprises, potentially, AMF as identified by 16:1(n-5). This marker was not separated to avoid mis-ID-ing bacterial lipoproteins (Nakayama et al., 2012), a possible error which would differentially affect soils from different landcover types as there are potentially more AMF in grasslands than in forests (see also supplementary Table S5).

### 2.9 Replication statement

**Table 1:** Overview to the scale at which the key parameters of this study were measured and their respective replication.

| Scale of inference | Scale at which the factor of interest is applied | Number of replicates at the appropriate scale |
| --- | --- | --- |
| **Rhizosphere respiration (plant root-soil-microbe continuum)** | Planted pot (d = 5.5 cm, V = 1.02 l) in individual respiration chambers (6.53l) | n=4 planted (15 plants per pot) per soil type |
| **Control soil respiration (soil-microbe)** | Unplanted pot (d = 5.5 cm, V = 1.02 l) in individual respiration chambers (6.53l) | n=4 unplanted per soil type |
| **Microbial biomass / PLFAs** | 5g / 10g soil per pot | n=4 planted + n=4 unplanted |
| **Photosynthesis (PS) and leaf area** | Leaf in ¼ l chamber | n=3 per pot, 4 pots per soil |

### 2.10 Statistical analysis

To identify whether the observed priming effects were significantly different between soil types and over time we first used a linear mixed model followed by t-tests using Satterthwaite’s method. Whether the total amount of primed C (All prime) and new SOC formation differed between soil types was each tested by a one-way ANOVA followed by post-hoc Tukey’s test. Variation in plant, soil and microbial parameters amongst soil types was analysed using principle component analysis (PCA) after Pearson (1901) on all parameters as measured on planted and unplanted soils, and where applicable the change in parameter between planted and unplanted was also included (e.g. delta microbial biomass C indicating the difference in microbial biomass C between planted and unplanted soils). To identify the potential drivers of RPE, from each ordination, we then used the two variables with the strongest eigenvalue of the first principal component to build linear regression models to predict each of the observed RPE (RPE1, RPE2, RPE3), the estimated total primed C (All prime) and newly formed soil organic carbon (New SOC). When two parameters had equal eigenvalues after three digits and were similar in ecological explanatory power, the one representing the more active elemental pool was chosen, that is soil mineral N rather than total soil N and proportion of plant-derived C in microbial biomass, rather than soil-derived C in microbial biomass and fungi in planted over fungi in unplanted soils (see also supplementary Tables S6A-D). Thereafter, we used stepwise deletion of terms based on Chi^2^ and model selection via AIC to reduce model complexity (Akaike, 1974). Models were validated inspecting plots of residuals vs fitted values, theoretical quantiles, standardized residuals vs fitted values and standardized residuals vs leverage (Cook’s distance). Statistical analysis was carried out using R 4.0.5 (R Core Team, 2021) with the additional packages car (Fox & Weisberg, 2019), cluster (Maechler et al., 2022), corrplot (Wei & Simko, 2021), ggfortify (Tang et al., 2016), ggpubr (Kassambara, 2020), lmerTest (Kuznetsova et al., 2017) and multcompView (Graves et al., 2015).

## 3 Results

### 3.1 RPE and SOC formation

Rhizosphere priming effects decreased in all soils over time (p< 0.001), including a shift from initially positive to finally negative priming in most soils (Fig. 1). The magnitude of priming was always greater in organic soils than in mineral soils, indicating a higher sensitivity of organic soils to RPE. For most time points, RPE was greater in sub-arctic soils than in their comparative Andean soils. The exceptions were positive priming in Andean Puna mineral soils at day 35, which was greater than in the boreal tundra mineral soils at that time, and by the negative priming in boreal forest soils at the later measurements, which was of similar magnitude than the negative priming in the Andean organic forest soils at that time (Fig.1). Only minor RPE was measured in the boreal mineral soils (BFM, BTM), while high and consistently positive RPE was measured in the organic tundra soil (BTO). The dynamics of RPE showed opposite trend in organic and mineral soils over time for both the arctic and the Andean soils. At the beginning of the experiment, RPE was higher in organic soils than in their mineral counterparts. At the end of the experiment, the opposite was observed: RPE was higher in the mineral soils as compared to their respective organic counterparts. One exception was the organic tundra heath soil (BTO), where RPE was always higher than in the mineral tundra heath soil (BTM). The sum of all priming effects (the integrated “All prime”) was close to zero as they changed from positive to negative priming (small plot in Fig.1), with the exceptions of overall negative RPE in organic Puna grassland soils (APO) and positive RPE in organic tundra heath soils (BTO). SOC formation was higher in organic than in mineral soils (Fig. 2A) and taken together with rhizosphere priming effects mostly caused net gain in soil C, apart from organic tundra soils (BTO), where net loss of C was observed as positive priming exceeded new SOC formation (Fig. 2B).

**Figure 1:**
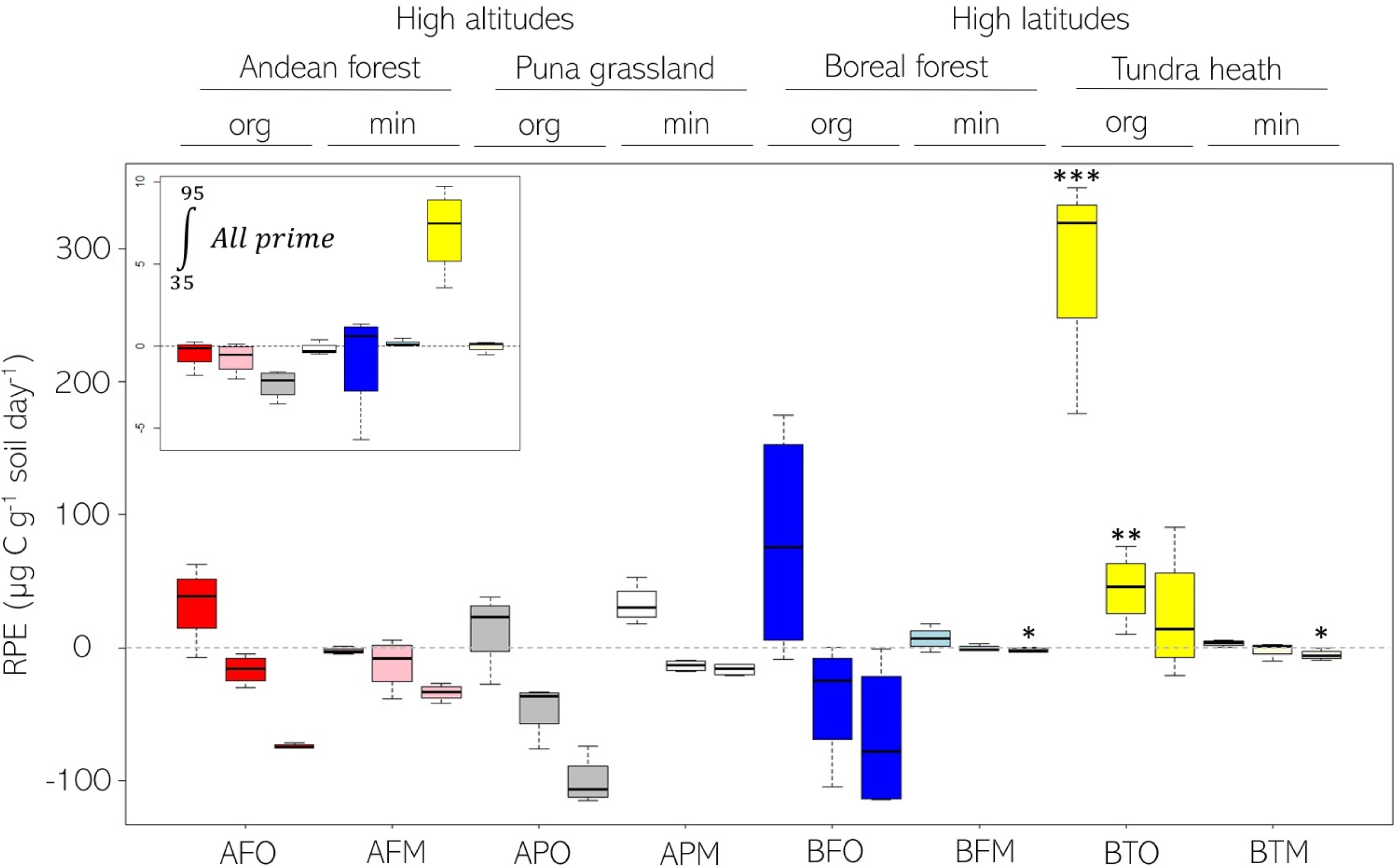
Rhizosphere priming effects. (RPE [μg C g-1 soil dwt day-1]) of eight treeline soils (AFO-BTM) each measured at three time points during the growing season (35, 80 and 95 days after sowing). Asterisk indicate differences between soil types at significance levels < 0.001 *** ≤ 0.01 ** ≤ 0.05 * according to linear mixed model followed by t-tests. Notably higher RPE in organic tundra soil (BTO) at the first two time points and in the mineral soils from the boreal subarctic (BFM & BTM) at the last measurement. Decreasing RPE over time in all soils. Small plot in top left corner shows total primed C [mg C g-1 soil] upscaled to the full experiment duration for each soil type (section 1.4).

**Figure 2:**
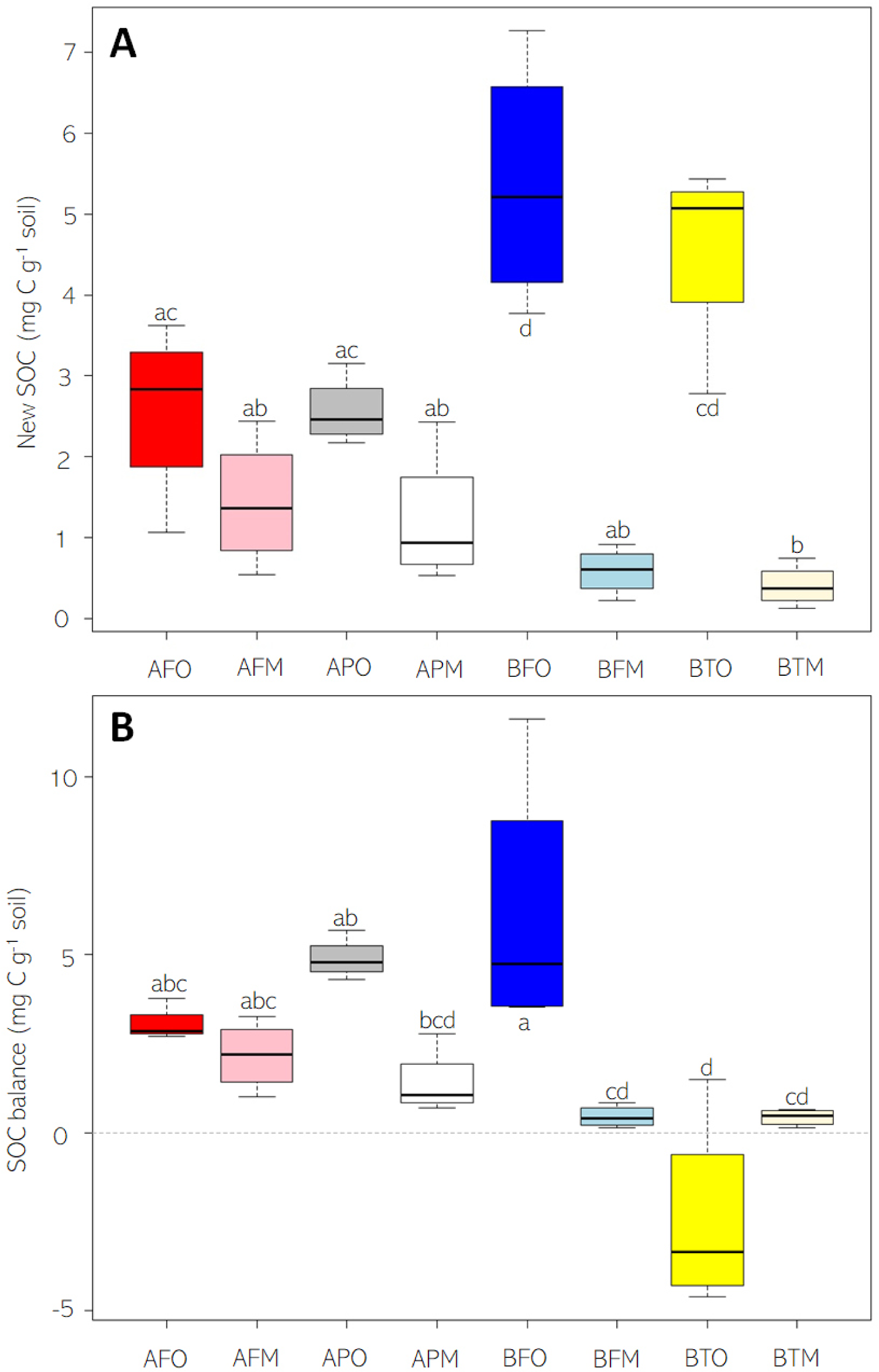
New soil organic carbon (SOC) formation. (A) and SOC balance (B) (mg C g soil-1) of eight treeline soils (AFO-BTM) at the end of a 95 days growing season. Net carbon loss (negative soil C balance in BFO soil). Letters indicate grouping after one-way ANOVA followed by Tukey’s post-hoc test.

### 3.2 Microbial carbon and nitrogen dynamics

Microbial biomass C and N were significantly different between soil types, with significant changes occurring following the introduction of live plants, quantified as the difference between unplanted and planted soils (Fig. 3). We observed mostly increases in microbial biomass C, with the greatest increase in organic boreal soils (BTO, BFO) and a slight decrease in Andean Puna grassland soils (APO, APM). For the Andean soils, the N gain exceeded C gains for all soils, resulting in lower microbial biomass C:N ratios in the planted soils. This was also observed for the boreal organic soils, while in the mineral soils, increase in microbial biomass C exceeded N gains, increasing C:N ratios. We also observed strong differences in the affinity of soil microbes to take up plant-derived carbon compounds (Fig. 3, Fig.5C), which was highest in mineral tundra soils (BTM) and lowest in organic Puna soils (APO).

**Figure 3:**
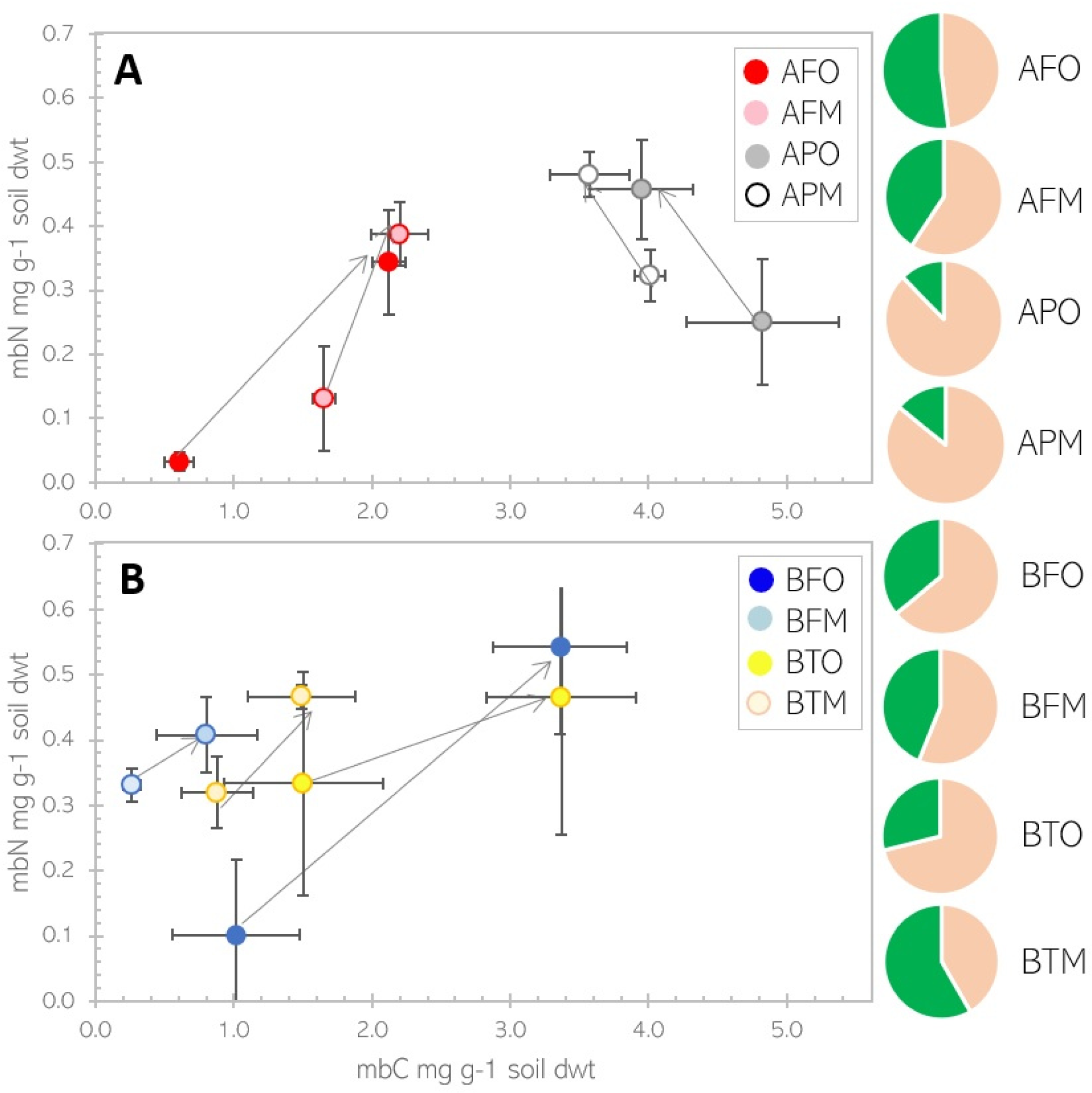
Microbial biomass carbon and nitrogen. (mg g-1 soil dry weight) in unplanted and planted treeline soils from Andean mountains (A) and Boreal subarctic (B). Arrows indicate change between unplanted and planted soils. Pie charts show the partitioning between soil (amber) and plant (green) derived microbial biomass carbon at the end of the experiment for each soil type.

### 3.3 Chemotaxonomic markers phospholipid-derived fatty acids (PLFAs)

Microbial community composition was determined as relative proportions of key functional groups of soil microbes as assigned to fatty acid biomarkers for fungi, actinomycetes, other gram positive and gram-negative bacteria and remaining unspecified PLFAs for n=4 replicates of each planted and unplanted soils for each soil type (Fig. 4). The size of the microbial community (total PLFAs) was not significantly different between planted and unplanted controls. Total concentrations of PLFAs were always higher in the organic soils compared to their mineral equivalents, with mineral soils having between ⅓ (Andean forest) to ⅒ (Boreal forest) of total biomarkers (p<0.001). Differences between soil types were mostly driven by gram-negative bacteria, actinomycetes and F:B ratios (Fig. 4, Fig. 5D). The boreal organic forest soils (BFO) had significantly more gram-negative bacteria than any other soil (p<0.001). Actinomycetes, like all other functional groups, were significantly higher in organic compared to mineral soils, but exceptionally low (p<0.001) in boreal mineral forest soils (BFM). The fungi-to-bacteria ratio (F:B) was significantly higher (p<0.001) in the organic tundra soils (BTM, Table 1).

**Figure 4:**
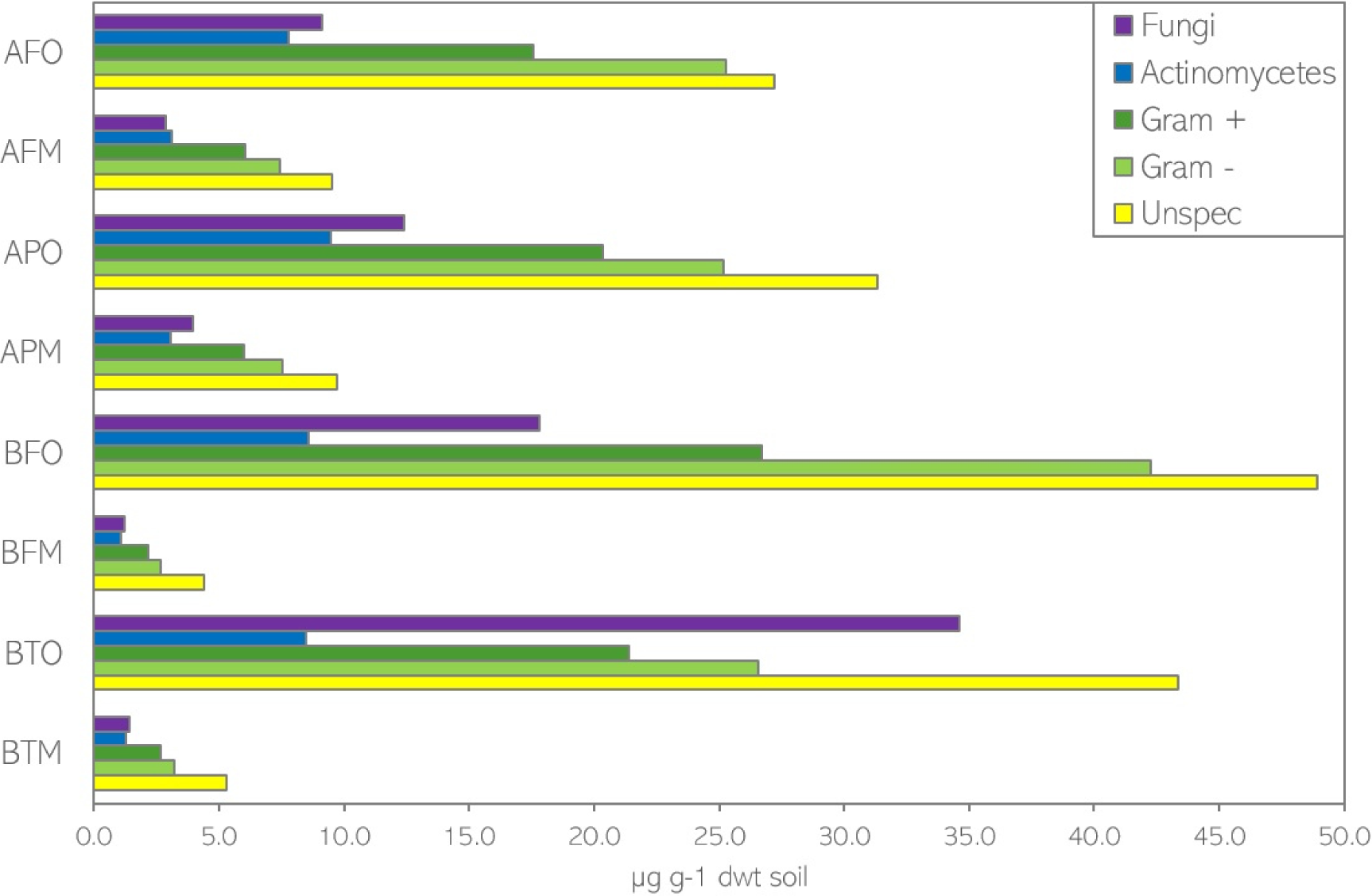
Total quantities of phospholipid fatty acid biomarkers. (PLFAs µg g-1 soil dry weight) assigned to functional groups of fungi, actinomycetes, gram positive and gram negative bacteria and unspecified soil microorganisms for eight treeline soils planted with *Cynodon dactylon*.

### 3.4 Plant growth depending on soil type

Plant growth and phenology strongly varied amongst the different soil types (Table 2, Fig. 5A). Leaf area index (LAI) was significantly higher for plants grown on organic boreal forest (BFO) soils at both measurements (p<0.001, p<0.01). Photosynthesis rates (PS) were only marginally different (p=0.06) for the plants in the different soil types at the beginning of the experiment, while later PS was highest for plants grown in Andean organic forest soils (AFO) and lowest for plants grown on their mineral counterparts (AFM). Root biomass was significantly lower (p< 0.001) in mineral tundra soils (BTM), while above ground plant biomass was significantly higher in the organic soils from Andean forest, Puna grassland and boreal forest (AFO, APO, BFO). Root-to-shoot ratios (R:S) were significantly lower in plants grown in the tundra soils, irrespective of soil horizon (p<0.01). Plants had in common that root nitrogen was below 1 %, while leaf nitrogen was above 1.5 %. Hence leaf tissue was enriched in N compared to root tissue, resulting in distinct C:N ratios (C:N leaves ≈ 25, C:N roots ≈ 60). Leaf C:N was significantly higher (p<0.01) for plants grown in mineral Puna grassland soils (APM) and lowest for plants grown in organic boreal forest soils (BFO). Root C:N was significantly higher (p<0.001) for plants grown in mineral tundra soils (BTM) and lowest for plants grown in organic boreal forest soils (BFO).

**Figure 5:**
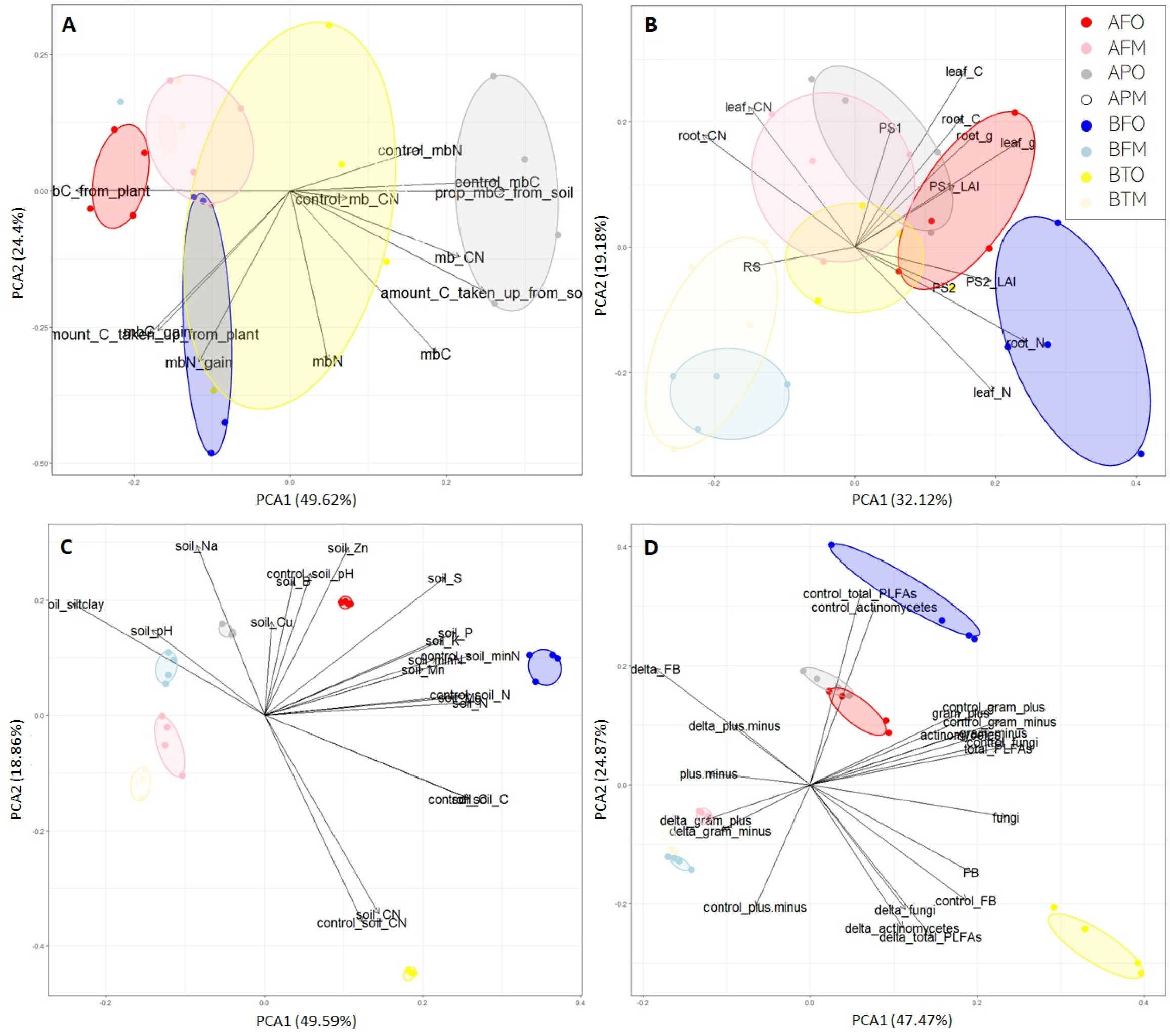
Ordination of chemical microbial parameters. (A), plant parameters (B), soil parameters (C) and taxonomic microbial parameter (D). Polygons group replicates of each soil type together. Full list of parameter definitions and PC loadings in supplementary material S6.

**Table 2:**
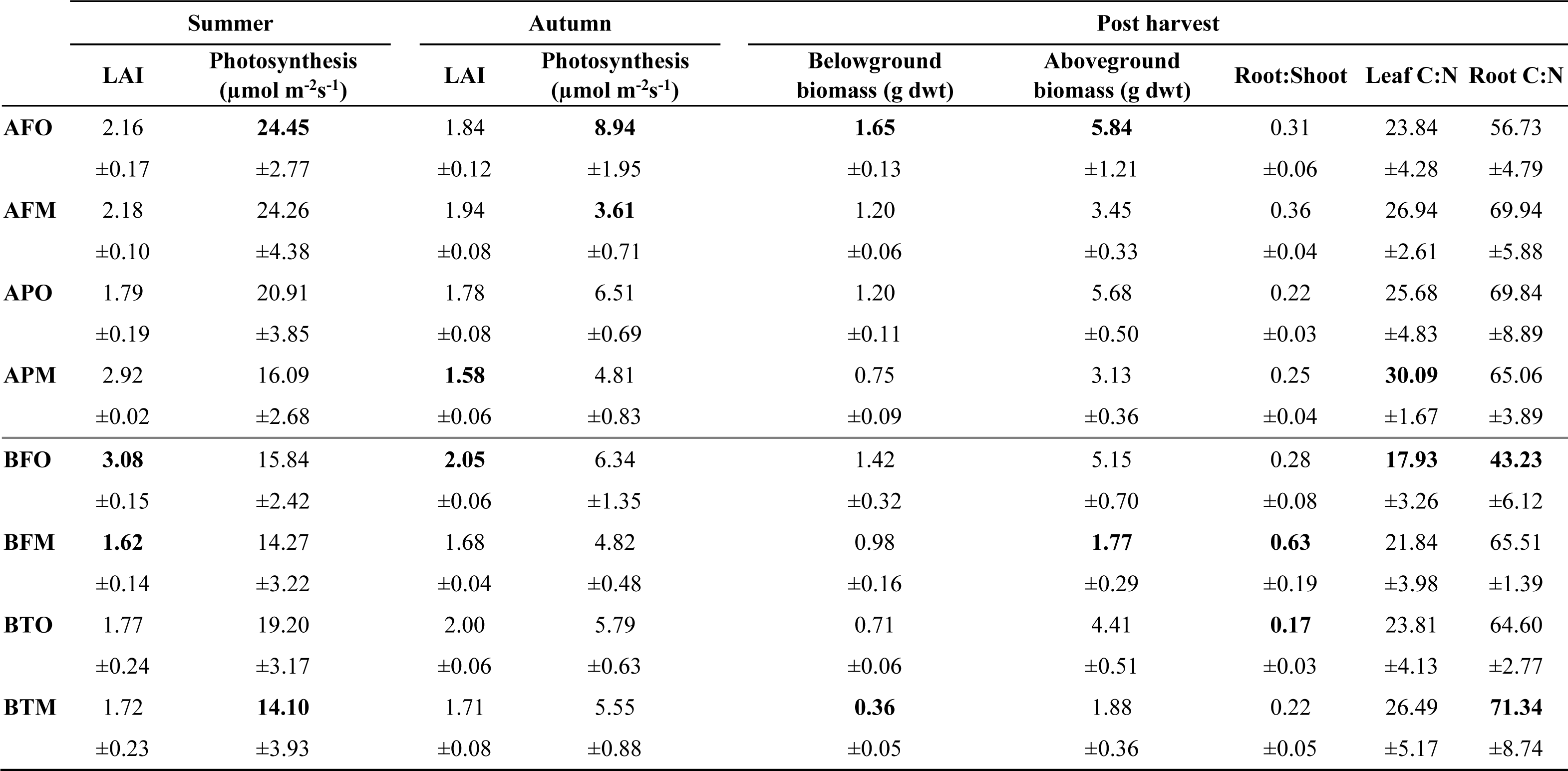
Plant parameter of *Cynodon dactylon* grown on eight different soil types. Leaf area index (LAI) and photosynthesis measured corresponding to RPE measurements, biomass and elemental composition determined after harvest. Maximum and minimum values highlighted in bold for each parameter.

### 3.5 Potential drivers of RPE and new SOC formation

First, principal component analysis (PCA) was used to disentangle the parameters in the plant, soil and microbial compartments that could best explain the observed differences between soil types (Fig.5). Soil and microbial parameters each explained around 70% of variance amongst soils with the first two principal components (PC), while plant parameters alone accounted for 50% variance with the two main PCs. Second, we combined the vectors with the strongest eigenvalue from each PCA in linear regression models to identify the parameters that best explain the observed RPE (Table 3). Soil carbon content was positively correlated with all priming estimates and new SOC formation, while soil mineral nitrogen was consistently negatively correlated with RPE estimates, but positively correlated with SOC formation. Microbial uptake of plant-derived C and fungi were mostly non-significant terms, but could not be removed from the model, indicating indirect relationships which cannot be captured by linear models alone. Moreover, root nitrogen content was positively correlated with RPE2 and the integrated All prime, as well as with SOC formation, while gram negative bacteria were negatively correlated with the (mostly negative) RPE3 towards the end of the experiment.

**Table 3:** Variables best explaining rhizosphere priming effects and new soil organic carbon formation. Coefficients of significant terms are given with significance codes < 0.001 ‘***’, ≤ 0.01 ‘**’, ≤ 0.05 ‘*’, ≤ 0.1 ‘.’ and non-significant but non-removable parameter indicated with ‘ns’. Parameters of model evaluation in last three columns (R^2^, p-value, AIC). All models started with the first two variables of the first principal component of each ordination (plant, soil, chemical microbes, taxonomic microbes) -> 8 variables used to build linear regression models, followed by stepwise deletion of terms based on Chi^2^ and model selection via AIC. Note that across the eight treeline soils, the first RPE is mostly positive (+), while RPE later is mostly negative (-) indicated by symbol in brackets (see also Fig.1 and 2).

|  | Plant<br>aboveground<br>biomass | Root<br>nitrogen | Soil<br>carbon | Soil<br>mineral<br>nitrogen<sup>a</sup> | Microbial<br>C uptake<br>from<br>plant | Microbial<br>biomass<br>carbon<sup>a</sup> | Fungi | Gram<br>negative<br>bacteria<sup>a</sup> | Multiple<br>R<sup>2</sup> | p-value | AIC (full +)<br>fitted model |
| --- | --- | --- | --- | --- | --- | --- | --- | --- | --- | --- | --- |
| **RPE1**<br>(+) |  |  | 1.07*** | -0.56** | ns | ns | ns |  | 0.61 | 0.0005 | (324.57)<br>320.37 |
| **RPE2**<br>(-) |  | 8070.7. | 0.37*** | -0.46*** | ns |  | -1.83. |  | 0.72 | 2.1e<sup>-5</sup> | (260.35)<br>255.56 |
| **RPE3**<br>(-) |  |  | 0.84*** | -0.28* | ns |  | ns | -5.67** | 0.73 | 1.9e<sup>-5</sup> | (267.84)<br>264.29 |
| **All**<br>**Prime** |  | ns | 0.04*** | -0.03** | ns |  | -0.2* |  | 0.84 | 6.2e<sup>-8</sup> | (112.52)<br>107.9 |
| **New**<br>**SOC** |  | 0.07. | 1.14e<sup>-2</sup>** | 7.32e<sup>-3</sup>* | ns |  | ns |  | 0.82 | 1.5e<sup>-7</sup> | (81.44)<br>79.68 |

<sup>a</sup> Parameter of unplanted control soil as suggested by PCA loadings (Fig. 5, supplementary material S6).

## 4 Discussion

Using natural abundance labelling and soils from high altitudinal and high latitudinal treelines, we observed that rhizosphere priming effects decreased during the plant growth season in all soils (Fig. 1). The impact of RPE on the overall soil C-balance in this experiment was minor in seven of the eight soils studied (Fig.2). This is in line with a previous study where priming effects were upscaled from field measurements and no indication was found that priming changes overall C-budgets (Cardinael et al. 2015). Similar observations were made by Schiedung et al. (2023) who evaluated priming effects along a 20-year chronosequence of land inversion in New Zealand to identify the dependence of priming effects on root-derived C in top soil and sub soils. Even though positive priming was reported, overall, carbon losses with priming never exceeded new root-derived carbon inputs. And also Yin et al. (2019), who studied rhizosphere priming effects and microbial biomass carbon dynamics of two wheat genotypes grown under two temperatures, found no net changes in soil carbon stocks as C losses caused by positive priming were counteracted by increased microbial growth/turnover. However, we did observe net soil C loss in one soil type, the sub-arctic organic tundra heath soil, which is in line with previous studies in this region that indicate a potential risk of net C losses in organic-rich subarctic soils caused by positive priming (Hartley et al., 2012; Keuper et al., 2020).

### 4.1 Magnitude of RPE depending on soil type

The magnitude of rhizosphere priming effects was always higher in organic compared to mineral soils (Fig. 1) and thus proportional to soil carbon content. (Table 1). In soils rich in organic matter, the potential for carbon loss from positive priming is naturally higher because the higher carbon density increases the spatial accessibility of carbon to soil microbes (Dungait et al. 2012). On the other hand, increased mineralization can increase the availability of nutrients, making them available for plant uptake. This can promote plant growth and productivity, which in turn increases atmospheric carbon removal via photosynthesis (Perveen et al., 2014; Bernard et al., 2022). It remains to be tested whether photosynthetic CO_2_-fiaxtion can compensate for CO_2_-loss from priming? The larger magnitude of RPE in organic soils suggests that microbes in organic rich soils have a higher reactivity to plant inputs than microbes in mineral soils, so organic soil C stocks may be more vulnerable to priming than mineral soil C stocks, which is in line with the findings of a similar study using switchgrass (de Graff et al., 2014) and a laboratory study (Salomé et al., 2009). In mineral soils, the magnitude of priming is naturally limited by lower carbon content, but also by smaller microbial communities, which have lower activities than larger microbial communities. The observed difference in RPE between organic and mineral soils could also be caused by higher SOM stability in deeper soils, where higher chemical recalcitrance and physio-chemical binding to minerals may protect carbon stocks from priming (Chen et al., 2019).

### 4.2 Direction of priming depending on plant growth stage and moment of measurement

Depending on the time of measurement, priming effects were either positive or negative. Both positive and negative priming are regularly reported in the literature and observed priming effects vary considerably in direction and magnitude amongst different studies (Bastida et al. 2019; Zheng et al. 2021). Therefore, without further long-term measurements, it remains impossible to predict the net impact of priming effects on the overall C balance at ecosystem scale (Qiao et al. 2014; Cardinael et al. 2015; Liang et al. 2018). In this experiment, rhizosphere priming became more negative over time and notably, this reversed the initially positive priming to eventually negative priming in most soils. Because of these dynamics, RPE did not lead to net C losses in most of the studied soils. However, this could have been concluded, had one measured RPE only once in September (RPE1). Long-term studies are therefore mandatory to provide realistic estimates of the real amplitude and impact of RPE on the C cycle. To better understand if this trend is having a large-scale impact on the studied ecosystems, we need to benchmark laboratory studies against ecosystem conditions embracing the complexity of plant-soil microbe interactions over time. Future studies could therefore evaluate the sensitivity of soil carbon to rhizosphere priming effects *in situ* and also trace the long-term fate of (labelled) plant inputs into microbial biomass and different soil fractions.

### 4.3 Carbon loss from subarctic organic tundra soils?

In this study, most soils showed a carbon balance in favour of C sequestration at the end of the experiment (Fig.2), yet net carbon loss from soil over time was observed for organic tundra soils, which is in line with other studies indicating carbon loss from positive priming in sub-arctic tundra heath and permafrost soils (Hartley et al., 2012; Wild et al., 2016; Hicks et al., 2020; Keuper et al., 2020). Positive priming effects are particularly concerning for ecosystems like the subarctic and the arctic, because these regions are particularly sensitive to climate change (Rantanen et al. 2022) and their soils have large C stocks, which could critically impact the global carbon cycle if mobilised to the atmosphere as CO_2_ (Su-Jong Jeong et al., 2018). On the other hand, recent studies show that warming arctic soils can also lead to significant negative priming effects (Verbrigghe at al. 2022), which has also been observed in substrate-addition experiments with arctic soils (Wild et al. 2023; Michel at al. 2023). Estimating the impact of priming effects on the ecosystem C balance in sub-arctic environments is therefore still critical, with some studies indicating that variables like the CO_2_-fixation potential of an increased aboveground biomass in a warming sub-arctic and restructuration processes in subsoils could prevent significant net C losses (Sistla et al., 2013).

### 4.4 Plant-inputs as primary biological drivers of RPE

The here observed rhizosphere priming effects followed a clear and uniform pattern according to plant growth stages through the growing season, decreasing over time in all soils (Fig.1). The experiment was conducted during peak plant growth in summer and leading into autumn. The observed seasonal pattern is in line with observations of seasonally fluctuating RPE in early spring, in summer and over longer study periods (Fu & Cheng, 2002; Shahzad et al. 2015; Zhang et al. 2017; Henneron et al. 2020, 2022). Such a seasonal pattern of rhizosphere priming reflects the metabolic activities of plants, notably high nutrient demand to sustain plant growth and decreasing photosynthesis and nutrient uptake from soil by the plant during senescence (Keiluveit et al. 2015; Girardin et al. 2016; Canarini et al. 2019). Plants directly sense environmental factors such as temperature and photoperiod, and adapt phenology, but also rhizosphere processes. At the time of the first RPE, plants were flourishing, reflected in high rates of photosynthesis and large LAI (Table 2). This likely had a two-fold effect in the rhizosphere: on the one hand, increased plant productivity also increases root exudation and hence the supply of fresh C to microbes (Kuzyakov & Cheng 2001; Jacoby et al. 2017; Guyonnet et al. 2018); on the other hand, increased plant productivity is likely also increasing the nutrient demand of the plant, detracting primary resources from microbes (Kuzyakov & Xu 2013; Canarini et al. 2019). One way of meeting an increased nutrients demand by the plant is to transfer C to soil microbes to enhance SOM-mineralisation, which is then manifested in positive priming and enhanced soil respiration (Cheng et al. 2012). On the other hand, root exudation is reduced with reduced plant photosynthetic activity, which also reduces the energy supply to microbes (Kuzyakov & Cheng 2001; Prudence et al. 2021). Negative rhizosphere priming occurred later in this experiment, where microbial degradation of SOM was reduced while photosynthesis decreased together with a decreased demand of nutrients by senescing plants and hence less inputs of labile C from plants to microbes (Prudence et al. 2021). In addition, the complexity and chemical composition of organic inputs change as plants age, with a shift from low molecular weight substances to a greater range of complex compounds in senescing roots and plant tissue, and / or a change in the C:N of these compounds (Cheng & Kuzyakov 2005; Zhang & Wang 2015). Direction and magnitude of priming depend partly on this change in chemical quality and quantity of inputs, and partly on the microbial functional capacity to utilise these different resources. Future studies could therefore measure plant exudation rates and exudate composition to improve the predictive power of the plant compartment on RPE (Fig. 5A, Table 3).

### 4.5 Microbial mediators of SOC turnover

The microbial community adapts to fluctuating plant nutrients demand and C inputs (Cotrufo et al. 2013; Dijkstra et al. 2013; Wang et al. 2016). Rates of SOM mineralization decrease when microbial demand for C and nutrients can be met by labile sources from exudates or other rhizodeposits through preferential substrate use (Blagodatsky et al. 2010; Blagodatskaya et al. 2011; Michel et al. 2023). Rates of SOM-mineralisation increase when microbes use the plant-derived C to mine nutrients from the soil (Cheng et al. 2005; Craine et al. 2007; Chen et al. 2014; Hicks et al. 2020). Substrate availability is a primary rate-limiting step of RPE and mediated by the plant (Mondini et al. 2006; Blagodatskaya et al. 2011; Gunina,et al. 2014; Jacoby et al. 2017). Microbial functional capacity and stoichiometric decomposition compose the second biological rate-limiting step of RPE (Rinnan & Baath 2009; Chen et al. 2014; Mooshammer et al. 2014). Our results further show a link between microbial uptake of plant-derived C and new soil organic matter formation (Fig. 3, Table 3). The uptake of plant-C into microbial biomass is likely an important step in soil formation, as microbial turnover generates necromass which can be rapidly adsorbed by soil minerals (Cotrufo et al. 2013; Gunina et al. 2014; Buckeridge et al. 2020). The incorporation of plant-derived C into microbial biomass was lowest in Andean Puna soils and highest in the mineral tundra soils (Fig. 3). Even though not always significant, fungal abundance was a permanent term in all models, and negatively correlated with the second and the integrated RPE (Table 3). Fungi can impact the C balance in two ways: on the one hand, fungi store more C in biomass than bacteria which can favour C-sequestration (Wilson et al. 2009; Drigo et al. 2010; He et al. 2020); on the other hand, fungi can increase C-loss from soil through release of SOM-degrading enzymes and breaking up soil minerals with their hyphae (Landeweert et al. 2001; Cheng et al. 2012; Finlay et al. 2020). The role of fungi in the C-cycle therewith remains unresolved and an important challenge for the future is to identify under which conditions fungi increase C-sequestration and when they do not. In addition to fungi, gram-negative bacteria were the microbial variables with the strongest eigenvalues in the PCA (Fig. 5D). Differences in the abundance of gram-positive or gram-negative bacteria could also impact the potential of new soil organic matter formation as they have different affinity to labile C-inputs and their specific cell wall compositions may differentially impact the recalcitrance and absorbance to soil minerals following cell turnover (Vollmer et al. 2008; Gunina et al. 2014; Enggrob et al. 2020).

### 4.6 RPE as a mechanism of plant-soil synchronisation

The plant-soil synchronization hypothesis (Swift, 1984; Myers et al. 1994) suggests that microbial mineralisation of SOM increases when plant demand is high and vice versa. In natural ecosystems, element cycles would thus be in equilibrium, as plant and microbial supply and demand are balanced over time. RPE fit this scheme as part of the synchronisation process, where microbes utilise the labile C provided through root exudation from plants to degrade SOM rich in nutrients, which are then available to plants for root uptake (Jacoby et al. 2017; Thirkell et al. 2019; Dellagi et al. 2020; Garcia et al. 2020). When plant nutrient demand is low, carbon supply can exceed microbial carbon demand and replenish the soil C stock. This manifests in negative priming effects, for example when the overall supply of nutrients in the rhizosphere exceeds plant and microbial demand (Dijkstra et al. 2013; Zhang et al. 2017). Negative priming can be further enhanced when plant productivity is reduced, like near the end of this experiment, where low photosynthesis and senescence were observed (Table 2). This continuous adjustment, which is manifested in more or less intense priming effects, could be further based on the principle of soils functioning as a bank, where plants and microbes dynamically debit and deposit into a continuous store of elements in the soil matrix (Fontaine & Barot, 2005; Perveen et al. 2014). On larger temporal and spatial scales, priming effects are hence not necessarily altering the overall ecosystem C balance, as they fluctuate according to situational plant and microbial supply and demand.

### 4.7 Challenges to upscaling RPE

For an integrative mechanistic understanding of positive and negative rhizosphere priming effects as ubiquitous processes in ecosystems, which continuously mediate carbon supply and nutrient demand between plants and microbes, it will be important to improve the methodological approaches to study RPE in presence of live plants and *in situ*. The detection and quantification of RPE can be biased by the time of sampling, as season and plant growth stage can have a strong impact on RPE (Fig.2). The here used natural abundance labelling has the advantage of providing continuous and realistic inputs of plant-derived carbon, while the system outputs, like plant nutrient, are equally unchanged and accounted for, which is not the case in laboratory soil incubations for example. The isotopic C partitioning which we did follows the usual rules in the field and is based on a two end-member mass balance approach (Subke et al. 2006; Kuzyakov 2011). However, it must be acknowledged that the natural variation in 13-C during plant growth and the relatively low labelling intensity induce uncertainty to the absolute values calculated using mass balance approach (supplementary material S3, S4). This approach does not allow to quantify the ‘primabilty’ of different SOC fractions (Paul et al. 2001; Bradford et al. 2008) and is valid only with the following assumptions: i) the isotopic signature of unlabelled pre-existent SOC determined by the 13-C measurement in the control soils is similar to the pre-existent SOC pool of the planted soils, where deviations are considered neglectable given the relative short duration of experiments (some months compared to 10-100 years of mean residence of soil C in topsoil) (Staddon 2004), ii) newly incorporated C is fresh C (exudates, rhizodeposition) and cannot be rapidly assimilated to the more evolved SOC (Blagodatskaya et al. 2011), iii) apparent priming caused by microbial biomass turnover is negligibly small as compared to the magnitude of real priming effects (Blagodatsky et al. 2010) and iv) microbial biomass per se remains relatively stable in unplanted soils, however minor 13-C fractionation occurs during baseline biomass turnover (Blagodatskaya et al. 2011; Lerch et al. 2011).

## 5. Conclusion

Whether climate change will amplify rhizosphere priming effects, and whether this will induce large-scale carbon losses from soils in vulnerable ecosystems remains an important question to predict the near future carbon balance of major ecosystems and estimate the feedback on atmospheric CO_2_ concentrations. Our results carry two important messages which indicate that we can currently not answer this question, because i) positive and negative priming effects are not mutually exclusive, rather omnipresent in ecosystems, and detection depends on the conditions of measurement and/or sampling time and ii) greenhouse and laboratory studies must be validated *in situ* to enable reliable ecological upscaling. For this purpose, methodological approaches to measuring RPE over time *in situ* under realistic conditions need to be significantly improved. To better understand the impact of treeline advance and higher plant productivity on the C balance, future studies could investigate the fate of plant C inputs in soils and belowground microbial and mesofauna communities, and quantify if enhanced SOM-mineralisation rates and soil CO_2_-emissions are compensated by enhanced plant photosynthesis and biomass production.

## Author Contributions statement

JM, SF and JW conceived the ideas and designed methodology; JM, SF, SR and CPC collected the data; JM analysed the data and led the writing of the manuscript. All authors contributed critically to the drafts and gave final approval for publication.

## Data availability

All data generated during the current study is presented in the manuscript and its supplementary files.

## Acknowledgments

We thank Robert Falcimagne, David Colosse and Ralf Boerger for technical support, Laurence Andanson and Christian Hussain for help with sample analysis and Iain Hartley for helpful comments and discussions. The financial support of the UK Natural Environmental Research Council (NE/L002434/1 and NE/S005137/1) and Campus France (mopga-short-0000000613) is greatly appreciated.

