## Supplemental material for "Vegetative stage and soil horizon determine direction and magnitude of rhizosphere priming effects in contrasting treeline soils"

Research Article

**S1: Table of soil micronutrient contents of eight treeline soils from Peru and Sweden.** CAL: Determination of phosphorus and potassium in calcium acetate lactate extract (VDLUFA I, A 6.2.1.1:2012), CAT: Determination of main and trace nutrients in calcium chloride/DTPA extract (VDLUFA I, A 6.4.1:2002).

| Sample ID | Land use | Soil horizon | Soil texture | pH (chalk) | P (mg in 100g)[1] | K (mg in 100g)[1] | Mg (mg in 100g)[1] | Cu (mg/kg) | Mn (mg/kg) | B (mg/kg) | Zn (mg/kg) | Na (mg/kg) | S (mg/kg) |
|---|---|---|---|---|---|---|---|---|---|---|---|---|---|
| | | | | $CaCl_2$ | CAL | CAL | $CaCl_2$ | CAT | CAT | CAT | CAT | CAT | CAT |
| **AFM** | Forest | Mineral | humus (4.1-8%), silty clay (silt > 50%) | 3.8 | 2 | 6 | 4 | 9.2 | 4 | < 0.10 | 12.2 | 15.6 | 4 |
| **AFO** | Forest | Organic | humus (8.1-15%), silty clay (silt > 50%) | 3.6 | 3 | 19 | 13 | 17.1 | 11.8 | 0.15 | 25.6 | 14.4 | 24.6 |
| **APM** | Grass-land | Mineral | humus (0-4%), silty clay (silt > 50%) | 4 | 1 | 5 | 3 | 13.7 | 3.3 | < 0.10 | 13.3 | 13.5 | 4 |
| **APO** | Grassl-and | Organic | humus (4.1-8%), silty clay (silt > 50%) | 3.9 | 2 | 11 | 5 | 18.4 | 3.4 | 0.12 | 20.8 | 13.6 | 11.7 |
| **BFM** | Forest | Mineral | humus (0-4%), very silty clay (silt > 50%) | 4.4 | 2 | 3 | 6 | 40.7 | 67.7 | < 0.10 | 27.8 | 9.3 | 5.1 |
| **BFO** | Forest | Organic | humus (> 30%), silt clay (-) | 4.6 | 5 | 15 | 12 | 24.7 | 162.3 | < 0.10 | 23.1 | 10.3 | 27.8 |
| **BTM** | Tundra | Mineral | humus (0-4%), very silty clay (silt > 50%) | 4.2 | 2 | 3 | 5 | 18.7 | 11.4 | < 0.10 | 13.1 | 9.2 | 5 |
| **BTO** | Tundra | Organic | humus (> 30%), silt clay (-) | 3.7 | 2 | 12 | 11 | 12.8 | 26.7 | 0.11 | 14.8 | 5.1 | 6 |

[1] Swedish organic: concentrations determined in soil solution ($H_20$) mg/100ml

**S2: Greenhouse air temperature and soil moisture contents**

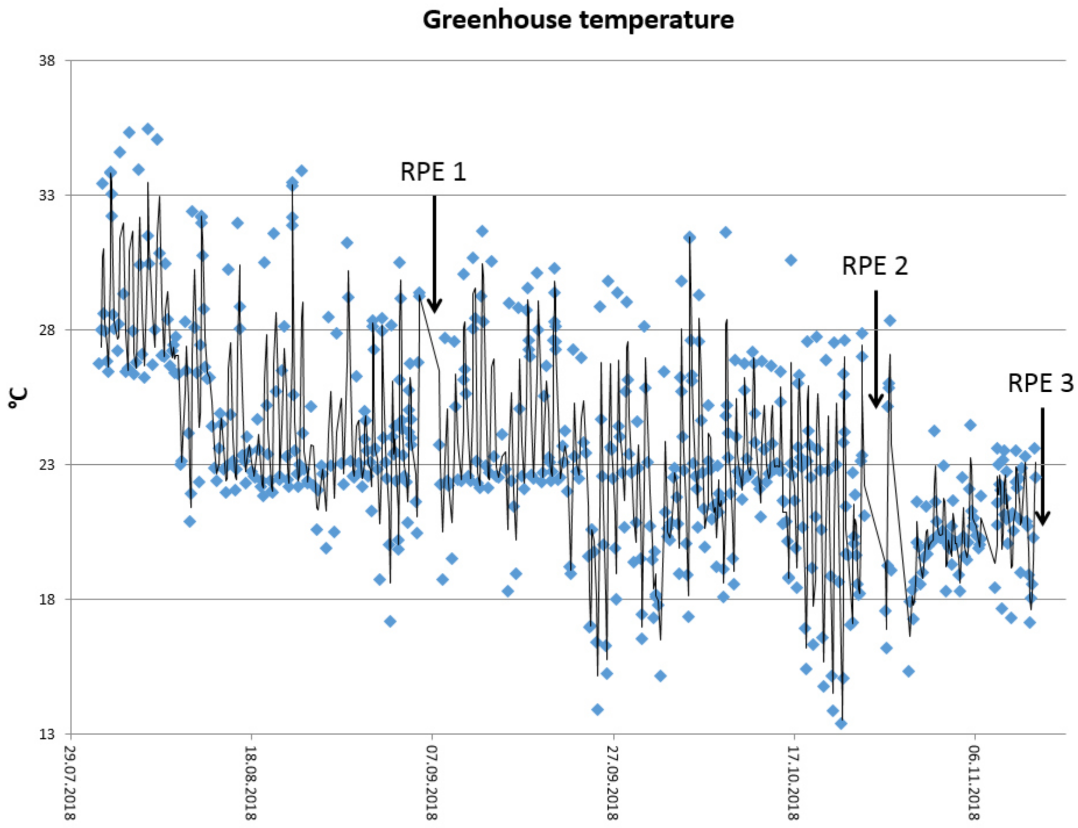

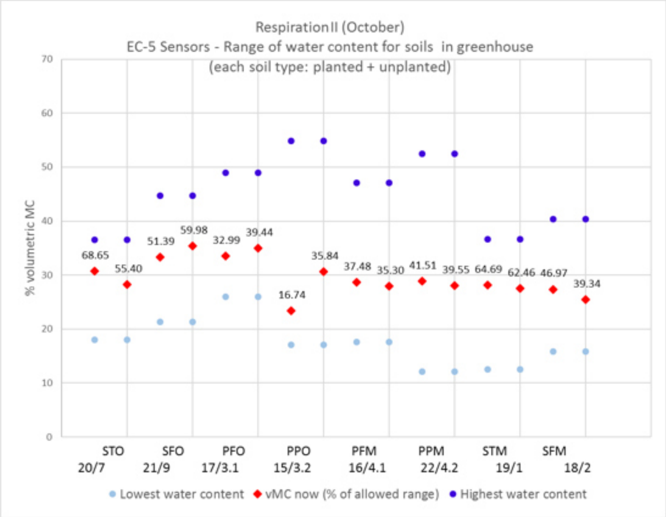

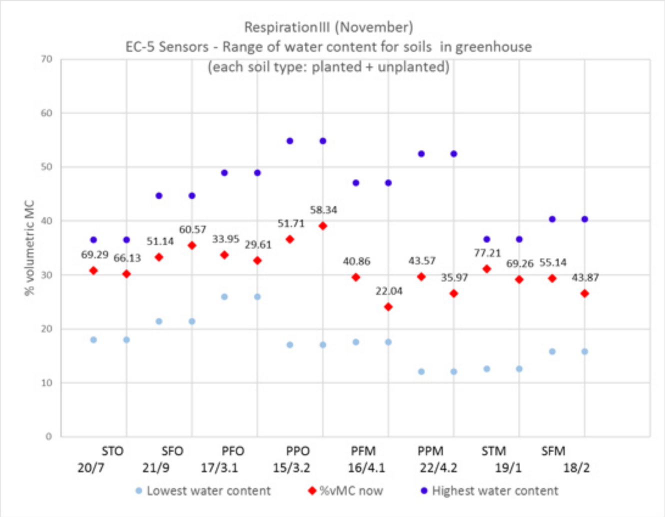

**S3: Table of 13C values for all compartments as measured during the experiment.**

| | Plant | | | | | | Soil | | | | |
|---|---|---|---|---|---|---|---|---|---|---|---|
| country x land cover x horizon | Leaves | | | | Roots | | Bulk | | | | Rhizosphere |
| | Respiration | | | Tissue | Respiration | Tissue | Respiration | | | Bulk | Respiration |
| **Soil ID** | **Sept** | **Oct** | **Nov** | **Dec** | **Nov** | **Dec** | **Sept** | **Oct** | **Nov** | **Dec** | **Nov** |
| **AFM1** | NA | -15.06 | -16.32 | -15.69 | -15.92 | -14.60 | | | | -25.41 | -17.68 |
| **AFM2** | -13.12 | -14.65 | -16.46 | -16.63 | -17.28 | -15.09 | | | | -25.47 | -17.64 |
| **AFM3** | -12.95 | -16.65 | | -15.44 | -16.15 | -14.72 | | | | -25.54 | -17.03 |
| **AFM4** | -13.38 | -15.57 | | -15.45 | -16.04 | -14.64 | | | | -25.28 | -19.56 |
| **AFM5** | | | | -15.58 | | | -21.67 | -23.58 | -19.20 | -25.59 | |
| **AFM6** | | | | -15.44 | | | -21.76 | -20.41 | -22.88 | -25.54 | |
| **AFM7** | | | | -16.28 | | | -21.22 | -24.01 | -23.07 | -25.64 | |
| **AFM8** | | | | -15.05 | | | NA | -23.93 | -23.05 | -25.65 | |
| **AFO1** | -11.82 | -15.45 | -16.08 | -15.58 | -16.71 | -14.85 | | | | -26.05 | -20.66 |
| **AFO2** | -12.42 | -15.71 | -16.46 | -15.44 | -15.20 | -14.58 | | | | -26.15 | -20.00 |
| **AFO3** | -12.55 | -14.98 | | -16.28 | -15.71 | -14.90 | | | | -26.06 | -15.74 |
| **AFO4** | -12.38 | -15.23 | | -15.05 | -16.15 | -14.88 | | | | -25.99 | -13.07 |
| **AFO5** | | | | | | | NA | -21.12 | -25.51 | -26.31 | |
| **AFO6** | | | | | | | -23.08 | -25.22 | -22.42 | -26.18 | |
| **AFO7** | | | | | | | NA | -26.30 | -25.02 | -26.11 | |
| **AFO8** | | | | | | | NA | -24.53 | -23.50 | -26.23 | |
| **APM1** | -11.47 | -15.83 | -16.11 | -15.18 | -17.03 | -14.87 | | | | -25.08 | -15.04 |
| **APM2** | -11.84 | -15.88 | -16.09 | -16.64 | -16.58 | -14.71 | | | | -25.03 | -16.10 |
| **APM3** | -11.54 | -16.28 | | -15.67 | -15.76 | -14.69 | | | | -25.11 | -18.73 |
| **APM4** | -11.42 | -16.37 | | -15.98 | -16.19 | -14.81 | | | | -24.83 | -15.07 |
| **APM5** | | | | | | | -20.22 | -20.90 | -24.53 | -25.23 | |
| **APM6** | | | | | | | -17.63 | -24.58 | -24.59 | -24.92 | |
| **APM7** | | | | | | | -19.04 | -23.31 | -23.20 | -25.31 | |
| **APM8** | | | | | | | -21.17 | -24.48 | -23.20 | -25.30 | |
| **APO1** | -13.14 | -16.23 | -16.91 | -15.62 | -15.89 | -14.70 | | | | -25.19 | -17.50 |
| **APO2** | -13.16 | -15.98 | -16.77 | -16.05 | -15.76 | -14.62 | | | | -25.14 | -17.70 |
| **APO3** | -12.35 | -16.30 | | -15.00 | -15.32 | -14.56 | | | | -25.21 | -18.79 |
| **APO4** | -13.53 | -15.61 | | -15.74 | -15.69 | -14.82 | | | | -25.19 | -18.25 |
| **APO5** | | | | | | | -18.06 | -24.04 | -25.02 | -25.34 | |
| **APO6** | | | | | | | -17.62 | -21.89 | -23.18 | -25.40 | |
| **APO7** | | | | | | | NA | -23.75 | -23.84 | -25.37 | |
| **APO8** | | | | | | | NA | -24.87 | -26.20 | -25.40 | |

| | | | | | | | | | | | |
|---|---|---|---|---|---|---|---|---|---|---|---|
| **BFM1** | -13.13 | -15.23 | -16.20 | -15.69 | -16.27 | -15.24 | | | | -27.00 | -19.51 |
| **BFM2** | -13.70 | -15.06 | -15.76 | -16.42 | -16.19 | -15.28 | | | | -27.15 | -19.89 |
| **BFM3** | -12.83 | -14.65 | | -16.36 | -16.25 | -15.20 | | | | -26.90 | -20.26 |
| **BFM4** | -14.06 | -16.65 | | -15.86 | -16.31 | -15.30 | | | | -26.74 | -18.38 |
| **BFM5** | | | | | | | -21.94 | -22.80 | -23.63 | -27.44 | |
| **BFM6** | | | | | | | -21.79 | -24.85 | -23.51 | -27.15 | |
| **BFM7** | | | | | | | -21.11 | -24.35 | -24.87 | -27.21 | |
| **BFM8** | | | | | | | -21.82 | -24.06 | -23.25 | -27.25 | |
| **BFO1** | -12.58 | -15.86 | -16.17 | -15.98 | -15.71 | -15.05 | | | | -28.68 | -23.38 |
| **BFO2** | -13.44 | -15.45 | -16.46 | -15.97 | -16.12 | -15.54 | | | | -28.81 | -24.06 |
| **BFO3** | -13.03 | -15.71 | | -15.94 | -16.06 | -15.22 | | | | -28.77 | -25.27 |
| **BFO4** | -12.43 | -14.98 | | -15.78 | -16.16 | -15.42 | | | | -28.73 | -24.12 |
| **BFO5** | | | | | | | -26.64 | -28.33 | -24.16 | -28.96 | |
| **BFO6** | | | | | | | -25.80 | -27.79 | -28.30 | -28.96 | |
| **BFO7** | | | | | | | -25.80 | -28.68 | -28.99 | -28.89 | |
| **BFO8** | | | | | | | -25.38 | -28.61 | -24.73 | -28.94 | |
| **BTM1** | -11.35 | -15.06 | -16.63 | -16.25 | -15.76 | -15.76 | | | | -26.03 | -19.68 |
| **BTM2** | -12.01 | -14.65 | -16.72 | -15.66 | -15.62 | -15.19 | | | | -25.80 | -17.27 |
| **BTM3** | -12.37 | -16.65 | | -14.99 | -15.96 | -14.60 | | | | -26.10 | -17.30 |
| **BTM4** | -12.14 | -15.57 | | -16.73 | -16.54 | -14.98 | | | | -25.96 | -17.23 |
| **BTM5** | | | | | | | -20.26 | -24.76 | -24.32 | -26.10 | |
| **BTM6** | | | | | | | -20.83 | -23.11 | -23.52 | -26.03 | |
| **BTM7** | | | | | | | -20.44 | -23.41 | -23.69 | -26.25 | |
| **BTM8** | | | | | | | -23.00 | -24.58 | -23.46 | -26.31 | |
| **BTO1** | -11.91 | -15.45 | -15.92 | -15.77 | -15.18 | -14.98 | | | | -27.11 | -24.12 |
| **BTO2** | -12.95 | -15.71 | -15.79 | -14.99 | -15.23 | -15.57 | | | | -27.04 | -21.22 |
| **BTO3** | -13.33 | -14.98 | | -16.24 | -15.69 | -15.23 | | | | -27.05 | -22.73 |
| **BTO4** | -13.98 | -15.23 | | -15.83 | -16.74 | -16.16 | | | | -27.06 | -23.62 |
| **BTO5** | | | | | | | -21.82 | -24.17 | -23.60 | -27.19 | |
| **BTO6** | | | | | | | NA | -26.49 | -26.29 | -27.20 | |
| **BTO7** | | | | | | | -20.92 | -24.88 | -24.86 | -27.13 | |
| **BTO8** | | | | | | | -21.64 | -24.78 | -24.52 | -27.21 | |

### S4: Uncertainty analysis

The RPE estimates obtained by the two end-member mass balance approach are a priori attached with uncertainty, because the calculations are based on several assumptions about the carbon fractions involved and their isotopic composition. Soil-released $CO_2$ is composed of carbon originating from the process of microbial mineralisation of different carbon fractions in the soil and therefor may vary over time and in its isotopic composition. The $^{13}C$ of the soil end-member can be represented by the $^{13}C$ of bulk SOM, but not all of the C in soil is actually available to microbes, so in the short term, the soil end-member is best represented by the $^{13}C$ of control soil respiration, where the $^{13}C$ represents the soil C fractions actually available to microbes. Therefore, in this study we assumed that the SOC pool accessible by soil microbes can be best inferred from control soil respiration and that isotopic fractionation and microbial biomass turnover are negligible (see section 3.6 in main manuscript). However, the isotopic signature of the SOC mineralised by microbes may change as a function of the quantity of 'old' SOC mineralised through RPE. For the plant end-member, organic inputs to soils, such as labile low molecular weight carbon like glucose and more complex molecules such as cellulose, also do not have the same isotopic composition (Collister et al. 1994) and thus provide a source of variability. A good proxy for the isotopic composition of plant inputs to soils can however be obtained by measuring the $^{13}C$ of plant tissues (roots and leaves) and additionally the $^{13}C$ of leaf and root respiration. The sum of these $^{13}C$ values allows to infer the full spectrum of $^{13}C$ in carbon compounds that plants can release to soil microbes. In this experiment, the measurable variation of endmember isotopic signature was up to ± 0.78 (plant) and ± 2.3 (soil), both depending on soil type (see supplementary Table S2).

To quantify how the choice of end-member $^{13}C$ can affect the quantitative estimate of RPE, we took the worst case scenario, which is variation in $^{13}C$ of soil and plant. For this purpose, we completed equation 3 from the main text with the various possible values for the $^{13}C$ of soil and plant. By these means various RPE values were calculated for each sample at each time point. We then calculated the average RPE (RPEav) and the standard deviation (stdv). We then calculated the coefficient of variation for each individual measurement of RPE. The uncertainty in RPE estimate was then expressed as:

$$\text{RPE uncertainty } = \left|\frac{\text{stdv}}{\text{RPE}_{\text{av}}}\right| x\ 100 \qquad \textit{(Eq S1)}$$

In difference to Cros et al. (2019) who used a theoretical variation of ±0.5‰, we included the actually measured $^{13}Cs$ of the soil end-member from both bulk SOC and soil $CO_2$-respiration, which is a greater variation (Table S3). Including the $^{13}C$ of SOM may overestimate the uncertainty because not all 'old' SOC is available to microbial mineralisation (otherwise SOC would never build up), yet at the same time it is exactly this fraction of SOM which is assumed to be mobilised by priming effects, which is why we included it in the uncertainty analysis. Future experiments could aim to narrow the range of carbon fractions in SOM, and their isotopic composition, which are mineralizable and which are not. In addition to the uncertainty derived from variable SOC fractions being mineralised by microbes, we took into consideration the variability of C inputs from plants. In this uncertainty analysis, we took all possible $^{13}C$s into account that were measured for the plant end-member during the full course of the experiment. This includes the measured $^{13}C$'s from leaf and root tissues and the measured $^{13}C$'s from their respiration. This defined a wide range of possible plant $^{13}C$ values (Table S3), assuming that carbon compounds released from the plant through rhizosphere processes can have any $^{13}C$ value ranging between the $^{13}C$ of structural components (tissue) and the $^{13}C$ of metabolic processes (carbon respired from leaves and roots). In sum, this revealed a vast potential uncertainty in some RPE estimates, which could be addressed in future studies to validate whether the current scientific consensus on using $^{13}C$ values from respiration provides the most accurate quantitative estimates of RPE and which proxies are most suited for the two-end member mass balance approach.

**Additional references uncertainty analysis:**

Collister JW, Rieley G, Stern B, Eglinton G, Fry B (1994) Compound-specific δ 13C analyses of leaf lipids from plants with differing carbon dioxide metabolisms. Organic Geochemistry 21(6–7) 619-627. https://doi.org/10.1016/0146-6380(94)90008-6.

Cros C, Alvarez G, Keuper F, Fontaine S (2019) A new experimental platform connecting the rhizosphere priming effect with CO2 fluxes of plant-soil systems. Soil Biology and Biochemistry 130: 12-22. https://doi.org/10.1016/j.soilbio.2018.11.022.

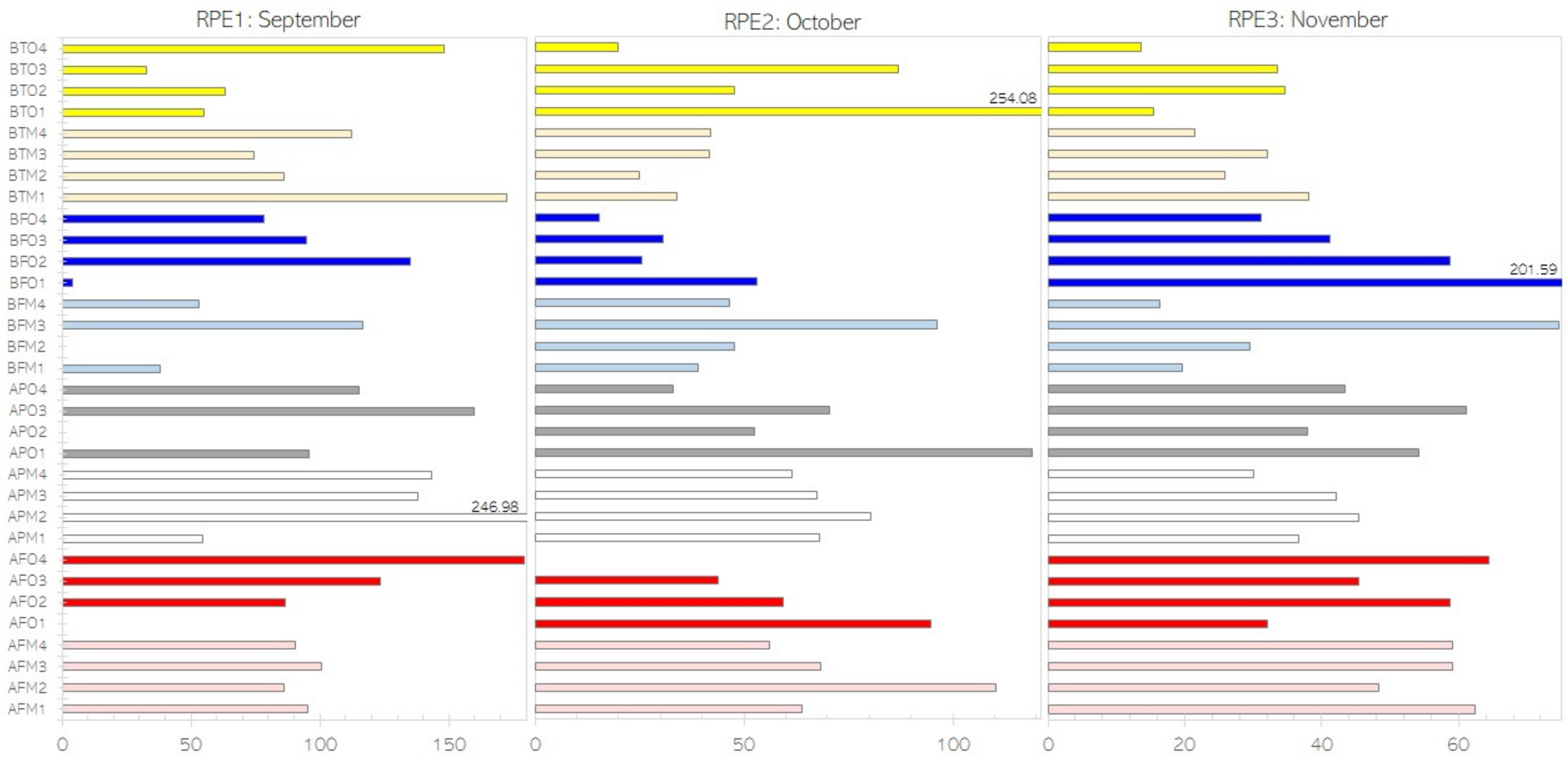

**Figure S4: Maximum potential uncertainty of RPE for each replicate of each soil type at each time point** (n=3-4), where each RPE is calculated using all possible $^{13}$C of plant inputs (root and leaf tissue and respiration) and two $^{13}$C of soil (bulk and respiration $^{13}$C).

**S5: Table of assignment of phospholipid-derived fatty acids (PLFAs) to functional groups.** In brackets markers which in this conservative assignment were included within unspecified PLFAs, although sometimes they are assigned differentially in the literature, as specified in brackets. Please see also section 1.7 main text.

| Fungi | Actino-mycetes | Gram + bacteria | Gram – bacteria | Unspecified PLFAs |
|---|---|---|---|---|
| **18:2** | 10Me-16:0 | 15:0i | 16:1(n-7) | 14:0i |
| **18:1(n-9)** | 10Me-17:0 | 15:0a | cy17:0 | 14:0 |
| | 10Me-18:0 | 16:0i | 18:1(n-7) | 15:1i |
| | | 17:0i | cy19:0 | 15:1a |
| | | 17:0a | (gram-) | 16:1(n-9) |
| (AMF) | | | | 16:1(n-5) |
| | | | (gram-) | 16:0 |
| | | | | 17:1i(n-8) |
| | | (bacteria) | (bacteria) | 17:0 |
| | | | | 18:0i |
| | | | (gram-) | 18:1(n-5) |
| | | | | 18:0 |
| | | | | 19:1 |

**S6: Tables of principal component (PC) loadings** for ordination of plant (A), soil (B), chemical microbial (C) and taxonomic microbial (D) parameters.

| **(A) Plant** | Parameter definition | Parameter unit | PC1 | PC2 |
|---|---|---|---|---|
| leaf_N | Leaf nitrogen content | mg g-1 dry weight | **0.312** | **-0.367** |
| leaf_C | Leaf carbon content | mg g-1 dry weight | 0.249 | **0.448** |
| leaf_CN | Leaf carbon to nitrogen ratio | unitless | -0.240 | **0.357** |
| **leaf_g** | Leaf biomass dry weight | gram | **0.370** | 0.268 |
| **root_N** | Root nitrogen content | mg g-1 dry weight | **0.386** | -0.241 |
| root_C | Root carbon content | mg g-1 dry weight | 0.240 | **0.326** |
| root_CN | Root carbon to nitrogen ratio | unitless | **-0.344** | 0.285 |
| root_g | Root biomass dry weight | gram | 0.272 | 0.286 |
| RS | Root to shoot ratio | unitless | -0.232 | -0.048 |
| PS1 | Photosynthesis at first RPE measurement | µmol m-2 s-3 | 0.083 | **0.305** |
| PS1_LAI | Leaf area index at first RPE measurement | unitless | 0.225 | 0.156 |
| PS2 | Photosynthesis at second RPE measurement | µmol m-2 s-3 | 0.202 | -0.102 |
| PS2_LAI | Leaf area index at second RPE measurement | unitless | **0.308** | -0.086 |
| Proportion of Variance | | | 0.321 | 0.192 |

| **(B) Soil** | Parameter definition | Parameter unit | PC1 | PC2 |
|---|---|---|---|---|
| **soil_C** | Soil carbon content (planted) | mg g-1 dry weight | **0.293** | -0.163 |
| control_soil_C | Soil carbon content (unplanted) | mg g-1 dry weight | 0.292 | -0.162 |
| soil_N | Soil nitrogen content (planted) | mg g-1 dry weight | 0.294 | 0.025 |
| control_soil_N | Soil nitrogen content (unplanted) | mg g-1 dry weight | 0.293 | 0.042 |
| control_soil_CN | Soil carbon to nitrogen ratio (unplanted) | unitless | 0.139 | -0.404 |
| soil_CN | Soil carbon to nitrogen ratio (planted) | unitless | 0.164 | -0.388 |
| soil_minN | Soil mineral nitrogen content (ammonium and nitrate) (planted) | µg g-1 dry weight | 0.245 | 0.110 |
| **control_soil_ minN** | Soil mineral nitrogen content (ammonium and nitrate) (unplanted) | µg g-1 dry weight | **0.294** | 0.118 |
| soil_pH | Soil pH (planted) | unitless | -0.160 | 0.166 |
| control_soil_pH | Soil pH (unplanted) | unitless | 0.066 | 0.278 |
| soil_siltclay | Soil silt and clay content | | -0.273 | 0.219 |
| soil_P | Soil phosphorous content[1] | mg in 100g dry weight | 0.272 | 0.162 |
| soil_K | Soil potassium content[1] | mg in 100g dry weight | 0.254 | 0.147 |
| soil_Mg | Soil magnesium content[1] | mg in 100g dry weight | 0.279 | 0.036 |
| soil_Cu | Soil cupper content[1] | mg kg -1 dry weight | 0.011 | 0.183 |
| soil_Mn | Soil manganese content[1] | mg kg -1 dry weight | 0.226 | 0.091 |
| soil_B | Soil boron content[1] | mg kg -1 dry weight | 0.041 | 0.264 |
| soil_Zn | Soil zinc content[1] | mg kg -1 dry weight | 0.120 | 0.329 |
| soil_Na | Soil sodium content[1] | mg kg -1 dry weight | -0.096 | 0.332 |
| soil_S | Soil sulphur content[1] | mg kg -1 dry weight | 0.258 | 0.270 |
| Proportion of Variance | | | 0.4969 | 0.1886 |

[1] Measured before planting.

| **(C) Chemical microbes** | Parameter definition | Parameter unit | PC1 | PC2 |
|---|---|---|---|---|
| mbC | Microbial biomass carbon (planted soil) | mg g-1 dry weight | 0.271 | -0.432 |
| **control_mbC** | Microbial biomass carbon (unplanted soil) | mg g-1 dry weight | **0.385** | 0.026 |
| mbC_gain | Difference in microbial biomass carbon between planted and unplanted soils | mg g-1 dry weight | -0.246 | -0.373 |
| **prop_mbC_from_plant** | Proportion of microbial biomass derived from plant inputs | unitless | **-0.398** | 0.002 |
| prop_mbC_from_soil | Proportion of microbial biomass derived from soil | unitless | 0.398 | -0.002 |
| amount_C_taken_up_ from_soil | Quantity of carbon taken up by microbes from soil | mg | 0.361 | -0.270 |
| amount_C_taken_up_ from_plant | Quantity of carbon taken up by microbes from plant inputs | mg | -0.261 | -0.382 |
| mbN | Microbial biomass nitrogen (planted soil) | mg g-1 dry weight | 0.070 | -0.452 |
| control_mbN | Microbial biomass nitrogen (unplanted soil) | mg g-1 dry weight | 0.242 | 0.110 |
| mbN_gain | Difference in microbial biomass nitrogen between planted and unplanted soils | mg g-1 dry weight | -0.169 | -0.455 |
| mb_CN | Microbial biomass carbon to nitrogen ratio (planted soil) | unitless | 0.315 | -0.176 |
| control_mb_CN | Microbial biomass carbon to nitrogen ratio (unplanted soil) | unitless | 0.105 | -0.020 |
| Proportion of Variance | | | 0.496 | 0.244 |

| **(D) Taxonomic microbes** | Parameter definition | Parameter unit | PC1 | PC2 |
|---|---|---|---|---|
| **fungi** | Biomarkers assigned to fungi (planted soil) | µg g-1 dry weight | **0.306** | -0.070 |
| control_fungi | Biomarkers assigned to fungi (unplanted soil) | µg g-1 dry weight | 0.301 | 0.095 |
| delta_fungi | Difference in fungal biomarkers between planted and unplanted soils | µg g-1 dry weight | 0.149 | **-0.273** |
| gram_plus | Biomarkers assigned to gram-positive bacteria (planted soil) | µg g-1 dry weight | 0.236 | 0.158 |
| control_gram_plus | Biomarkers assigned to gram-positive bacteria (unplanted soil) | µg g-1 dry weight | **0.287** | 0.172 |
| delta_gram_plus | Difference in gram-positive bacteria biomarkers between planted and unplanted soils | µg g-1 dry weight | -0.162 | -0.076 |
| gram_minus | Biomarkers assigned to gram-negative bacteria (planted soil) | µg g-1 dry weight | 0.286 | 0.114 |
| **control_gram_minus** | Biomarkers assigned to gram-negative bacteria (unplanted soil) | µg g-1 dry weight | **0.296** | 0.139 |
| delta_gram_minus | Difference in gram-negative bacteria biomarkers between planted and unplanted soils | µg g-1 dry weight | -0.140 | -0.100 |
| FB | Fungal to bacteria ratio (planted soils) | unitless | 0.251 | -0.191 |
| control_FB | Fungal to bacteria ratio (unplanted soils) | unitless | 0.243 | -0.253 |
| delta_FB | Difference in fungal to bacteria ratio between planted and unplanted soils | unitless | -0.242 | 0.254 |
| actinomycetes | Biomarkers assigned to actinomycetes (planted soil) | µg g-1 dry weight | 0.235 | 0.111 |
| control_actinomycetes | Biomarkers assigned to actinomycetes (unplanted soil) | µg g-1 dry weight | 0.101 | **0.390** |
| delta_actinomycetes | Difference in actinomycetes biomarkers between planted and unplanted soils | µg g-1 dry weight | 0.144 | **-0.312** |

| | | | | |
|---|---|---|---|---|
| total_PLFAs | Sum of phospholipid fatty acids extracted from planted soils | μg g-1 dry weight | **0.293** | 0.081 |
| control_total_PLFAs | Sum of phospholipid fatty acids extracted from unplanted soils | μg g-1 dry weight | 0.078 | **0.419** |
| delta_total_PLFAs | Difference in phospholipid fatty acids extracted from planted and unplanted soils | μg g-1 dry weight | 0.189 | **-0.332** |
| plus.minus | Ratio of gram-positive to gram-negative bacteria (planted soils) | unitless | -0.157 | 0.026 |
| control_plus.minus | Ratio of gram-positive to gram-negative bacteria (unplanted soils) | unitless | -0.086 | -0.267 |
| delta_plus.minus | Difference in ratio of gram-positive to gram-negative bacteria between planted and unplanted soils | unitless | -0.117 | 0.130 |
| Proportion of Variance | | | 0.475 | 0.249 |

### S7: Alternative RPE graphs

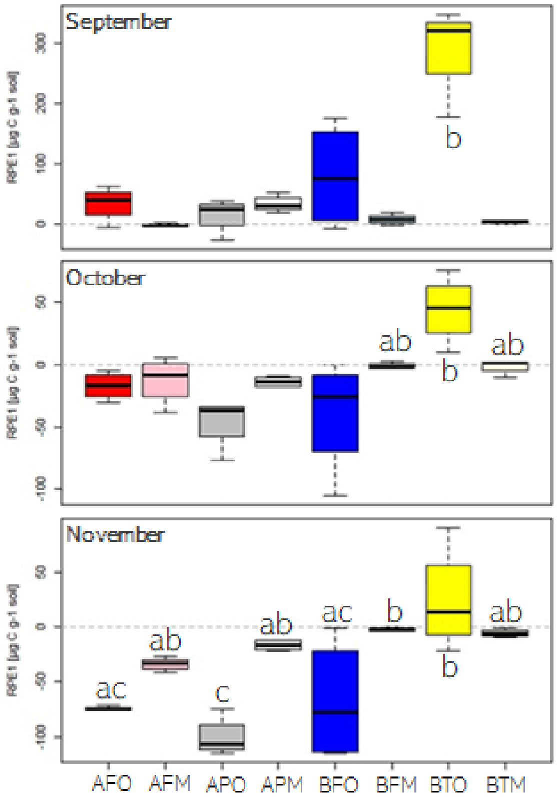

Figure S7.1: RPE [µg C $g^{-1}$ soil] measured over eight soil types three times during the growing season. Letters indicate differences between soil types following ANOVA and post-hoc Tukey's test for each time point individually ($p < 0.001$ for all three months).

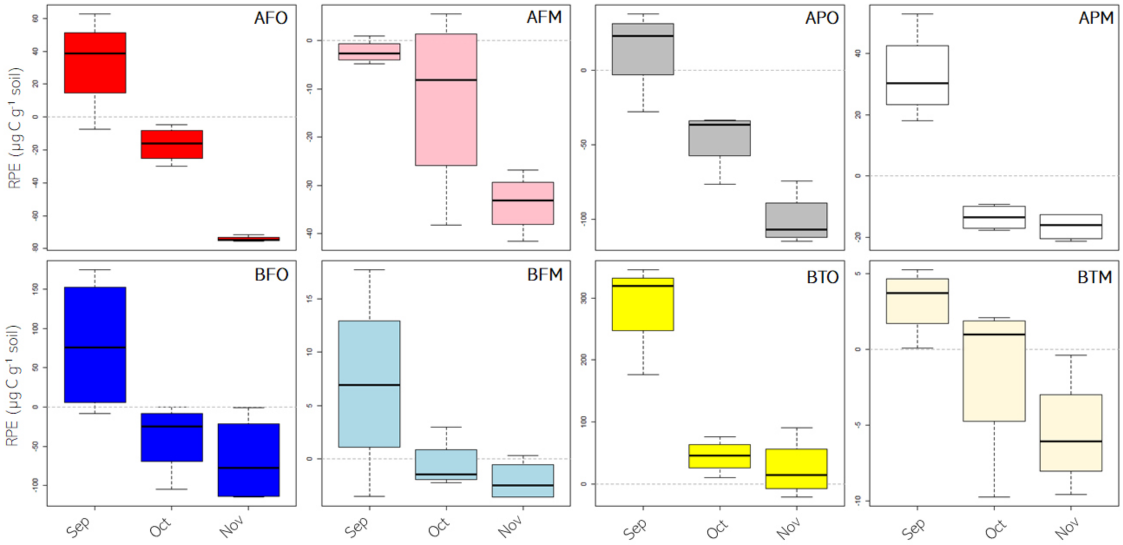

Figure S7.2: RPE [µg C $g^{-1}$ soil] measured over eight soil types three times during the growing season.

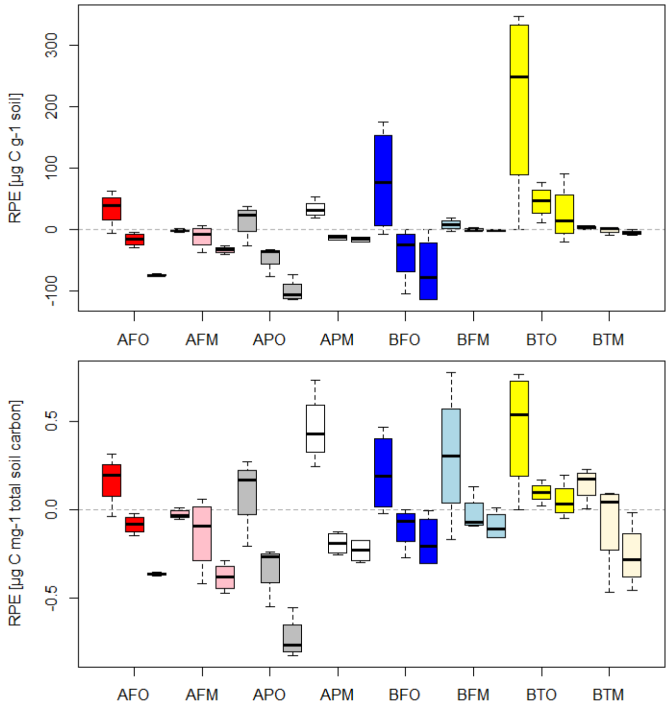

Figure S7.3: RPE measured over eight soil types three times during the growing season. Top panel: RPE expressed per gram soil dry weight [µg C $g^{-1}$ soil]. Bottom panel: RPE expressed per milli gram total soil carbon [µg C $mg^{-1}$ total soil carbon].
